# Molecular classification of zebrafish retinal ganglion cells links genes to cell types to behavior

**DOI:** 10.1101/2020.07.29.226050

**Authors:** Yvonne Kölsch, Joshua Hahn, Anna Sappington, Manuel Stemmer, António M. Fernandes, Thomas O. Helmbrecht, Shriya Lele, Salwan Butrus, Eva Laurell, Irene Arnold-Ammer, Karthik Shekhar, Joshua R. Sanes, Herwig Baier

## Abstract

Retinal ganglion cells (RGCs) form an array of feature detectors, which convey visual information to central brain regions. Characterizing RGC diversity is required to understand the logic of the underlying functional segregation. Using single-cell transcriptomics, we systematically classified RGCs in adult and larval zebrafish, thereby identifying marker genes for at least 33 stable and transient cell types. We used this dataset to engineer transgenic driver lines, enabling experimental access to specific RGC types. Strikingly, expression of one or few transcription factors often predicts dendrite morphologies and axonal projections to specific tectal layers and extratectal targets. *In vivo* calcium imaging revealed that molecularly defined RGCs exhibit highly specific functional tuning. Finally, chemogenetic ablation of *eomesa*^*+*^ RGCs, which comprise melanopsin-expressing types with projections to a small subset of central targets, selectively impaired phototaxis. Together, our study establishes a framework for systematically studying the functional architecture of the visual system.

## Introduction

Understanding how the brain regulates behavior requires targeted genetic access to subpopulations of neurons, so they can be characterized and manipulated. Retinal ganglion cells (RGCs) are the sole output neurons of the eye and therefore represent a bottleneck through which all visual information flows as it is transmitted to visual centers of the brain. RGCs can be subdivided into several dozen types based on anatomical, physiological and molecular characteristics (Baden et al., 2016; Bae et al., 2018; Dacey, 1994; Peng et al., 2019; Robles et al., 2014; Tran et al., 2019; Yan et al., 2020). Their selective tuning to certain visual features, such as luminance transitions, edges, chromaticity or direction of motion, arises from inputs provided by specific subsets of retinal interneurons in the inner plexiform layer (**Figure 1A**), as well as intrinsic properties (Sanes and Masland, 2015). In many cases, distinct RGC functional classes project axons to different visual brain centers, which in turn are associated with specific perceptual and behavioral functions (Dhande et al., 2015; Kramer et al., 2019; Martersteck et al., 2017; Nikolaou et al., 2012; Robles et al., 2013, 2014; Seabrook et al., 2017; Semmelhack et al., 2014; Temizer et al., 2015). Both the individual synaptic connectivities of RGCs and their biophysical characteristics are determined to a large extent by their cell type-specific gene expression profiles. Owing to the key role they play in early visual processing, RGCs are prime targets for a functional dissection of visually guided behavior.

**Figure 1.**
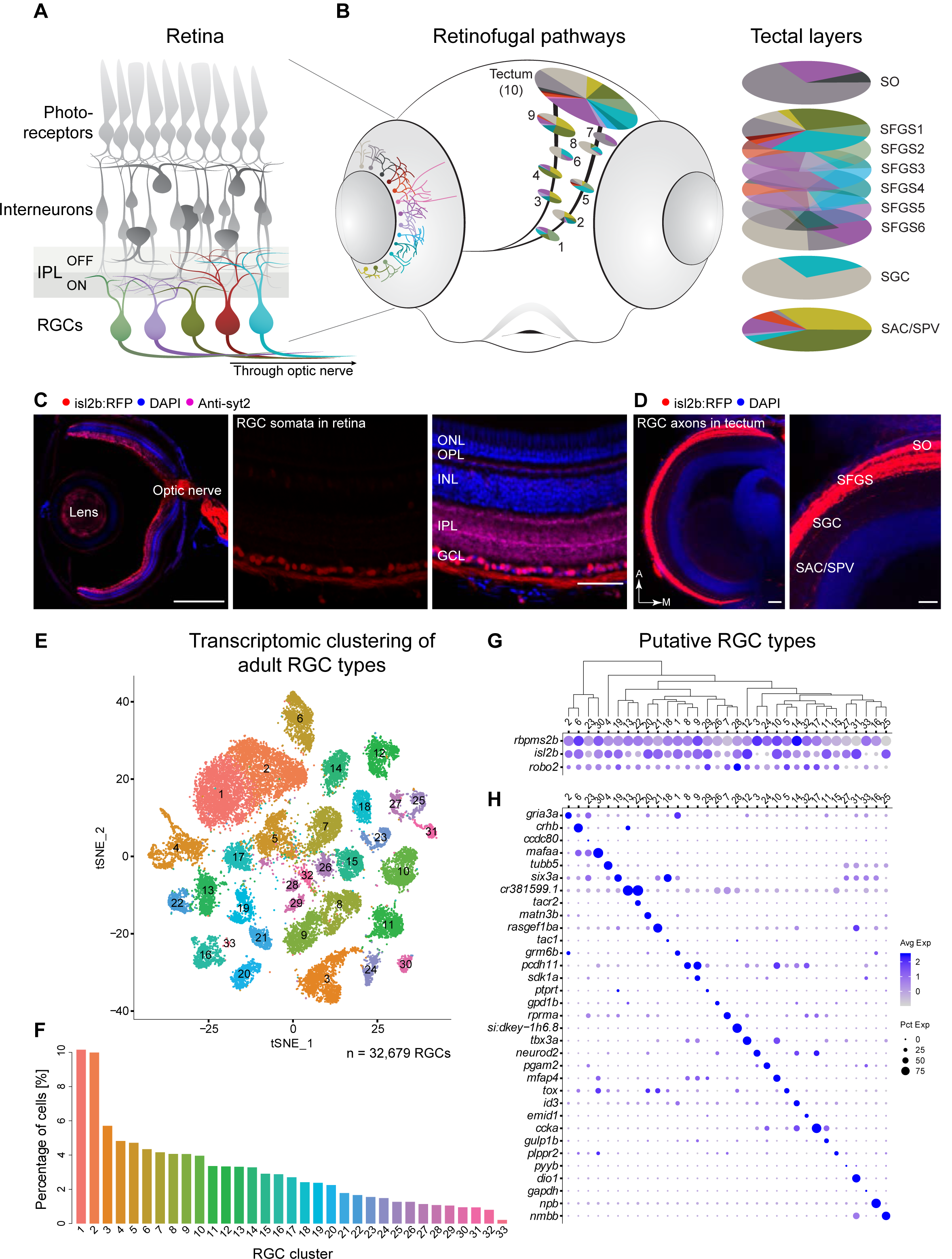
Single cell transcriptomics defines an adult zebrafish RGC catalog comprising 33 molecularly distinct clusters. **(A)** Sketch of the zebrafish retina. RGCs, the innermost retinal neurons, transmit visual information to the rest of the brain through the optic nerve. Unique patterns of dendritic stratification in the inner plexiform layer (IPL) enables distinct RGC types (colors) to receive presynaptic input from specific interneuron types, rendering individual RGC types sensitive to distinct visual features. **(B)** *Left:* The zebrafish ‘Retinal Projectome’ (Robles et al., 2014). Morphological RGC types are defined by stereotyped combinations of dendritic profiles in the retina and axonal projections to retinorecipient nuclei, named arborization fields (AFs) 1-9 and tectum. *Right:* Within the tectum, the main innervation site, RGC axons form nine laminar innervation domains: SO (stratum opticum), SFGS (stratum fibrosum et griseum superficiale) 1-6, SGC (stratum griseum centrale), and the boundary between SAC/SPV (stratum album centrale/ stratum periventriculare). Each AF or tectal lamina is innervated by a unique set of RGC morphotypes, depicted by colors. **(C)** *Tg(isl2b:tagRFP)* labels RGCs. *Left:* Section of an adult eye immunostained for RFP, synaptotagmin (syt2) as a neuropil reference and DAPI counterstain of somata. *Middle:* A magnified retinal section highlighting RFP-labeled RGCs. *Right:* Overlay of all markers in the retinal section. ONL, outer nuclear layer; OPL, outer plexiform layer; INL, inner nuclear layer; IPL, inner plexiform layer; GCL, ganglion cell layer. Scale bars, 500 µm (*left*), 50 µm *(middle and right)*. **(D)** *Left:* Confocal plane covering the RFP-immunostained adult *Tg(isl2b:tagRFP)* tectum with DAPI counterstain. *Right*: Magnified area. A, anterior; M, medial; SO, stratum opticum; SFGS, stratum fibrosum et griseum superficiale; SGC, stratum griseum centrale; SAC/SPV, boundary between stratum album centrale and stratum periventriculare. Scale bar, 100 µm. **(E)** t-distributed stochastic neighbor embedding (tSNE) visualization of 33 transcriptional clusters (colors) of 32,679 adult zebrafish RGCs (points). Clusters are numbered in the order of decreasing relative frequency. **(F)** Relative frequency (y-axis) of adult RGC clusters (x-axis), ordered from highest to lowest. Clusters are colored as in E. **(G)** Dotplot showing the expression patterns of three classical zebrafish RGC markers (rows) across adult clusters (columns). The area of each circle depicts the percentage of cells expressing the gene, and the color depicts the z-scored expression in cells with non-zero transcripts. Clusters are ordered based on their global transcriptional relatedness depicted using a dendrogram (top), computed using hierarchical clustering. **(H)** Dotplot showing expression patterns of markers (rows) that are selectively enriched in adult RGC clusters (columns). Column ordering and expression depiction as in G.

Recent studies have used high-throughput single cell RNA sequencing (scRNA-seq) to generate comprehensive molecular taxonomies of RGC types in mice (Rheaume et al., 2018; Tran et al., 2019), non-human primates (Peng et al., 2019) and humans (Yan et al., 2020). However, it has remained challenging to associate individual types with their postsynaptic targets and the behaviors their activation elicits. The zebrafish is an attractive model system to address this gap for several reasons (Orger, 2016; Orger and de Polavieja, 2017): (1) Despite having diverged from mammals >400 million years ago, major aspects of retinal architecture and development are conserved in zebrafish (Sanes and Zipursky, 2010). (2) Zebrafish are suitable for rapid and efficient transgenic manipulation at large scale, which enables genetic changes to be connected to phenotypes (Baier and Scott, 2009). (3) Their visual system develops rapidly and a diverse visual behavioral repertoire is seen by 5 days post fertilization (dpf) (Fleisch and Neuhauss, 2006; Orger, 2016; Orger and de Polavieja, 2017). (4) Zebrafish larvae are transparent, allowing for imaging of structure and function *in vivo* (Baier and Scott, 2009). (5) RGC types have been comprehensively catalogued in zebrafish, based on their morphologies and projection patterns to their retinorecipient targets (Kunst et al., 2019; Robles et al., 2013, 2014). These studies have revealed that RGCs send their axons to ten different layers in the tectum, as well as nine extratectal arborization fields (AFs), which are numbered according to their position along the optic tract AF1 through 9 (**Figure 1B**). These AFs are neuropil areas of brain nuclei in the hypothalamus/preoptic area, thalamus and pretectum, which are highly conserved among vertebrates and subserve a wide array of visual functions, such as the detection of optic flow, the photo-entrainment of circadian rhythms, prey recognition, and phototaxis. According to their behavioral function, each of the ten tectal layers and the nine AFs receives a distinct combination of feature-selective RGC inputs. Definitive analysis of these visual processing channels, however, has been hampered by a dearth of molecular markers, which are required to link cellular morphologies and physiological profiles of individual RGC types to the behaviors they subserve.

Here, we used scRNA-seq to comprehensively classify zebrafish RGCs based on their transcriptomes. We generated and compared the cell type catalogs of adult and larval RGCs. Nearly two thirds of larval RGCs exhibited molecular profiles that correspond to their adult counterparts, suggesting that a substantial proportion of the RGC repertoire is established at early larval stages. The remaining third were RGCs largely committed to specific adult fates, but were still in the process of transitioning to their mature state. One small cluster of cells, which persisted into adulthood, comprised uncommitted RGCs, reflecting continued neurogenesis in the teleost retina. Using CRISPR-Cas9 genome engineering and intersectional strategies, we established transgenic lines to gain exclusive genetic access to several molecularly defined RGC types. Anatomical characterization revealed that different molecular types project in stereotyped patterns to visual brain nuclei. We functionally characterized a small set of RGC types defined by selective expression of the T-box transcription factor *eomesa* using *in vivo* calcium imaging. These *eomesa*^*+*^ RGCs predominantly respond to increases in ambient luminance and also specifically express melanopsin-coding genes. Selective chemogenetic ablation experiments showed that *eomesa*^*+*^ RGCs regulate phototaxis. Overall, our study provides an inroad for systematically investigating the development, structure, function, and behavioral contributions of specific cell types in the vertebrate visual system.

## Results

### Single cell transcriptional profiling generates a molecular taxonomy of RGCs

We isolated RFP-labeled RGCs from adult (4-6 months old) transgenic *Tg(isl2b:tagRFP)* zebrafish (**Figure 1C-D**) and profiled them using droplet-based scRNA-seq (Zheng et al., 2017). Through computational analysis of transcriptional profiles, we separated major cell classes based on the expression of canonical markers. RGCs comprised 67% of cells, with the remainder including rods, Müller glia, amacrine cells, bipolar cells and endothelial cells (**Figure S1A**). We recovered a total of 32,679 high-quality single RGC transcriptomes with an average of 2,570 transcripts and 1,188 genes per cell. RGC transcriptomes were batch-corrected, and analyzed using dimensionality reduction and clustering (**STAR Methods)**, yielding 33 transcriptionally distinct clusters with frequencies ranging from 0.2% to 10.1% (**Figures 1 E-F, S1B-D**). All clusters expressed one or more of the pan-RGC markers *rbpms2b, isl2b* and *robo2* (**Figure 1G**). Most clusters could be uniquely labeled based on selective expression of a single gene, but in a few cases unique labeling required two genes (**Figures 1H, S1E, Table S1**). Transcriptomically distinct clusters likely represent individual cell types or groups of few cell types.

### RGC clusters are distinguished by expression of transcription factors and secreted molecules

Transcription factors (TFs) play essential roles in determination of cell identity (Fishell and Heintz, 2013). We therefore surveyed TF expression in our transcriptomic data. From a database of 1,524 TF-encoding genes, we found 186 that were expressed in >30% of cells in at least one RGC cluster (**STAR Methods**). About 10% (n=17) of these TFs were broadly expressed in most types. The majority (n=169) showed highly variable expression profiles, which were often restricted to one or few clusters (**Figures 2A-B, Table S2**). Restricted TFs could be further subdivided into two groups based on their degree of cluster-specific expression: 42% (n=71) exhibited multi-type expression in >20% of clusters and included the well-studied RGC-specific POU class 4 homeobox genes *pou4f1* and *pou4f2*, as well as *pou6f2, barhl2* and *ebf3a*. The remaining TFs (n=98) were expressed in 1-6 clusters each and included *eomesa/tbr2, mafaa, tbr1b, foxp2, satb1, satb2* and *mef2cb*.

**Figure 2.**
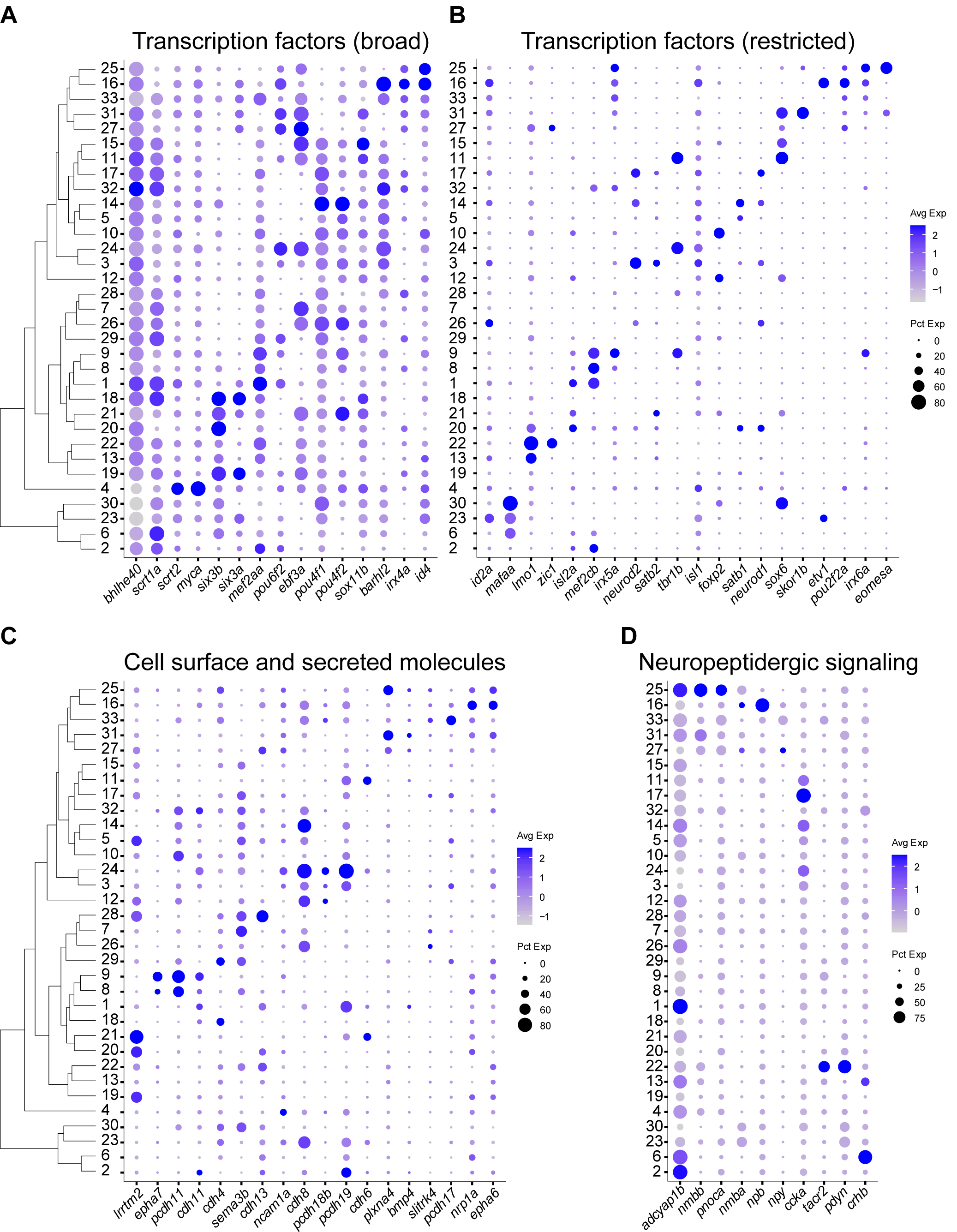
Variably expressed transcription factors, cell surface and secreted molecules, and neuropeptides across adult RGC types. **(A)-(B)** Dotplots highlighting examples of variably expressed TFs in adult RGC clusters, subdivided into broad (A), and restricted (B) categories. Representation as in **Figure 1G**. The full list is provided in **Table S2**. **(C)** Dotplot highlighting key cell surface and secreted molecules selectively expressed in adult RGC clusters. **(D)** Dotplot highlighting neuropeptides selectively expressed in adult RGC clusters.

In addition to TFs, we also identified variably expressed cell-surface recognition molecules, secreted guidance molecules, and neuropeptides (**Figures 2C-D, Table S2**). These included genes encoding ephrin receptors, *epha6* and *epha7*, which have been implicated in retinotopic axon guidance (Kita et al., 2015), and type II cadherins, known to regulate dendritic targeting in the IPL (Duan et al., 2014, 2018). Among neuropeptides, selectively expressed genes included *nmbb, npb, ccka* and *tacr2*. Overall, our data indicate that the molecular identity of individual RGC types is largely determined by the combinatorial expression of TFs, cell-surface molecules and neuropeptides.

### scRNA-seq highlights diversification of RGCs at the larval stage

To survey the molecular diversity of larval RGCs, we profiled RFP-positive cells from 5 dpf *Tg(isl2b:tagRFP)* fish (**Figures S2A-C**). We recovered 11,046 RGCs comprising 90.4% of all cells collected (**Figure S2D**). Our data represent a 10-fold enrichment and >3X coverage over their baseline frequency of 9% with ∼3,500 RGCs in the larval retina (Robles et al., 2014; Zimmermann et al., 2018). Larval RGCs were analyzed using the pipeline that had been applied to the adult data, yielding 29 clusters (**STAR Methods, Figures 3A and S2E-I**). Of these, 23 corresponded to adult RGC types by several criteria, and could be identified using specific markers (**Figure 3B, C, S2J**). We also observed that a statistically significant proportion of transcription factors (p < 10^−132^; hypergeometric test), recognition molecules (p < 10^−44^) and neuropeptides (p < 10^−13^) specifically expressed in the adult clusters retained their specificity among larval clusters (**STAR Methods, Figure S3A-D**).

**Figure 3.**
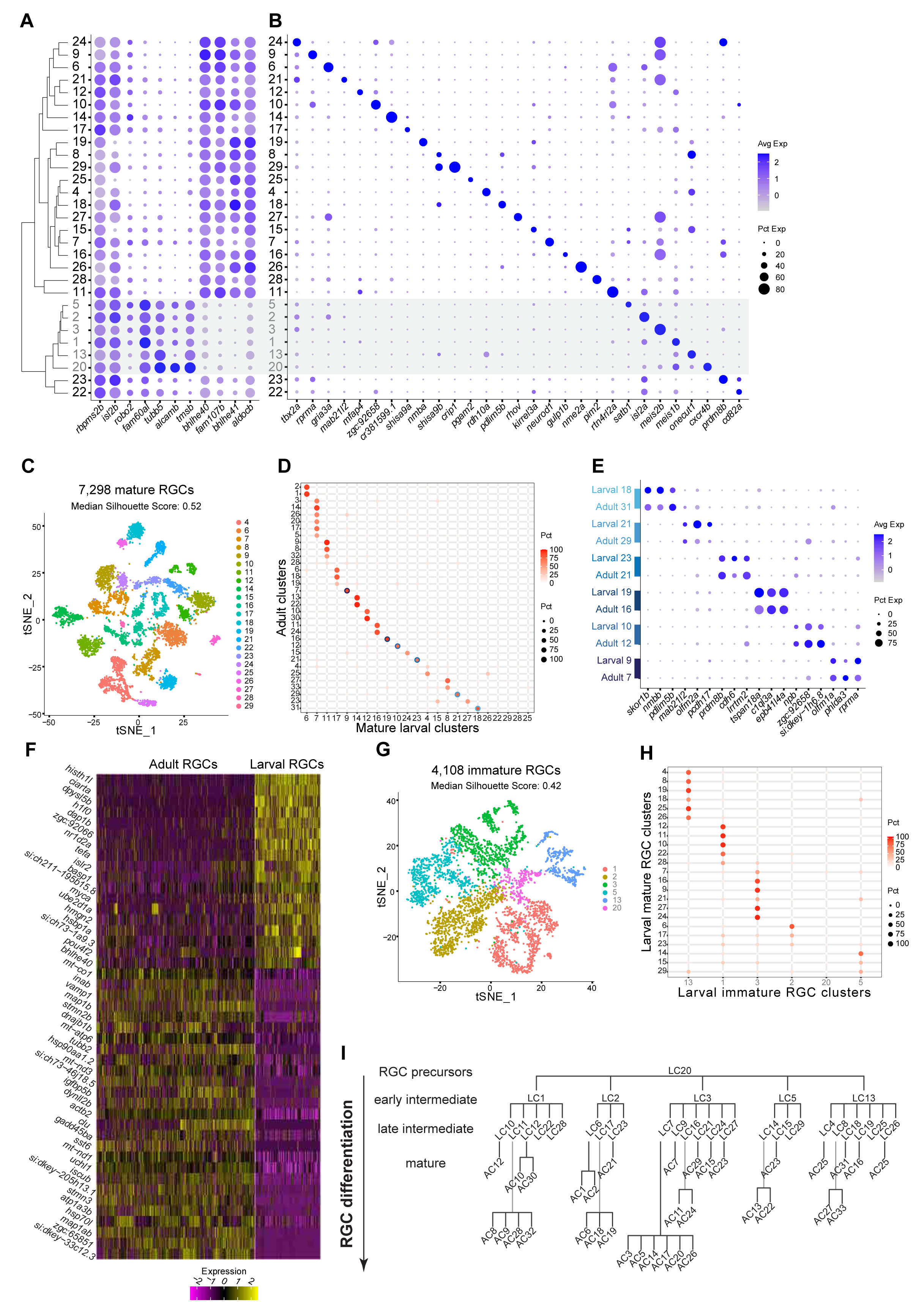
Molecular classification of larval RGCs and their transcriptomic correspondence with adult RGC types. **(A)** Global transcriptional relatedness (dendrogram, left) of larval RGC clusters (rows) identifies two groups, corresponding to mature and immature RGCs (grey cluster labels and shaded horizontal bar). The dotplot highlights expression of pan-RGC markers *rbpms2b, isl2b* and *robo2* as well as the top differentially expressed genes (n=8) between immature and mature RGC clusters. **(B)** Expression patterns of markers (columns) that are selectively enriched in larval RGC clusters (rows), ordered as in panel A. **(C)** tSNE visualization of 23 mature RGC clusters (colors) comprising 7,298 cells (points). The cluster IDs were used to compute a silhouette score for each cell based on its tSNE coordinates. The score ranges from -1 to 1, with higher values indicating tighter cluster boundaries. The median value of this score (top) quantifies the discreteness of the clusters. We note that the tSNE coordinates were not used to define the clusters. **(D)** Transcriptional correspondence between adult and mature larval RGC clusters. Circles and colors indicate the proportion of cells in an adult RGC cluster (row) mapped to a corresponding mature larval cluster (column) by the xgboost algorithm trained on mature larval RGCs. Each row is normalized to sum to 100%. Blue circles highlight six instances of a 1:1 corresponding pair of adult and larval clusters, which are separately analyzed in panels E-G. **(E)** Dotplot showing shared patterns of gene expression between the six 1:1 cluster pairings selected from the classification model (blue circles in panel D). Colored bars (left) indicate matching cluster pairs. **(F)** Heatmap showing differentially expressed genes (rows) between adult and larval stages identified from the six 1:1 matching clusters shown in panel D. Columns correspond to individual RGCs grouped by age. Values are row-wise z-scored gene expression values. **(G)** tSNE visualization of 6 immature RGC clusters (colors) comprising 4,108 cells (points). The median silhouette score was computed as in C. **(H)** Transcriptional correspondence between mature (rows) and immature (columns) larval RGC clusters using an xgboost classifier trained on immature larval RGCs. Representation as in panel D. **(I)** Model of RGC type diversification. Larval clusters (LCs) and adult clusters (ACs) are arranged by their transcriptional correspondences shown in D and H. RGC precursors give rise to immature (early intermediate) RGC clusters and mature (late intermediate) larval clusters, which further diversify into mature adult clusters.

Supervised classification analysis demonstrated that six of these clusters mapped to single adult clusters in a 1:1 fashion, likely representing types whose diversification is complete at the larval stage (**Figure 3D**). We verified these mappings based on the shared expression of cluster-specific markers at both stages (**Figure 3E**). Twelve additional larval clusters mapped to a few adult types each (see below). Four small clusters, each comprising less than 2.3% of RGCs, could not be matched with any adult counterparts, perhaps because they matured into types too rare to be identified in our atlas. We found 57 differentially expressed genes between the 1:1 matched larval and adult RGC types associated with global maturational changes (**STAR Methods, Figure 3F**). The most notable pattern was that genes with sequence-specific DNA binding activity were higher at the larval stage. Examples include *histh1l, nr1d2a, h3f3b*.*1, bhlhe40*, and *ciarta* (Howe et al., 2012). Genes whose expression levels increased with maturation were associated with mature neuronal function, such as *snap25a, vamp1, nefma* and *igfbp5b* (Howe et al., 2012). In addition to these global changes, we also found maturational changes that were type-specific (**Figure S3F**). Thus, our results reveal a substantial level of RGC molecular diversification in larval zebrafish that undergoes further refinement as the fish ages.

### Some larval clusters represent immature yet committed RGC types

In addition to the 23 larval RGC types with clear relationship to adult types, 6 transcriptionally proximate clusters (36% of larval RGCs) were defined by a distinct gene expression signature (**Figure 3A**). Notably, these genes were associated with early RGC development, such as the axon outgrowth gene *alcamb*, known to regulate early RGC axon outgrowth in zebrafish (Diekmann and Stuermer, 2009) as well as *tubb5* and *tmsb*, which are associated with cytoskeletal rearrangement (Ngo et al., 2014; Roth et al., 1999). We therefore reasoned that these clusters may represent immature larval RGCs.

We compared the two groups of clusters, finding three distinctions that support their division into immature and mature larval RGC types. First, genes defining mature larval RGCs (e.g. *bhlhe40, bhlhe41* and *fam107b*) were expressed widely across adult RGC clusters. In contrast, immature larval RGC genes such as *alcamb, tubb5* and *tmsb* were expressed in a single adult RGC cluster (C4), comprising only 4.8% of adult RGCs (**Figure S3E**). Second, clusters of mature larval RGCs were more transcriptionally distinct compared to immature larval RGCs, as judged by their tighter separation in the reduced dimensional space, quantified using the silhouette score (**STAR Methods, Figure 3C, G**). Third, supervised classification analysis showed that five out of the six immature clusters were transcriptionally related to nonoverlapping sets of 3-7 mature larval clusters (**Figure 3H**). Interestingly, top differentially expressed genes for these immature RGC clusters included *satb1, isl2a, meis2b, meis1b* and *onecut1* (**Figure 3B**), which are TFs that continue to be selectively expressed among mature larval RGCs and adult RGCs (**Figures 2A-B, S3A-B**). The observation that the mature types to which immature types mapped were mutually exclusive supports the idea that these five RGC clusters represent committed precursors restricted to specific fates that are gradually refined. Finally, the sixth immature cluster (C20), comprising 2.3% of all larval RGCs, had the highest expression of *alcamb, tubb5*, and *tmsb*, and exhibited transcriptional correspondence to adult cluster 4 (data not shown). The unique markers that distinguished this cluster included *fgf8*, a factor involved in initiation of RGC differentiation (Martinez-Morales et al., 2005), and *cxcr4b*, a receptor involved in early axon migration (Li, 2005). We propose that C20 might represent uncommitted precursors.

Attempts to resolve instances of “multi-mapping” of larval to mature types by searching for further substructure in larval clusters were unsuccessful (**Figure S3G-I**). While this may reflect insufficient resolution in the larval cells due to low RNA capture, it is also possible that the diversification of these cells is truly incomplete at the molecular level. Taken together, these results are consistent with a diversification model in which transcriptionally distinct immature RGCs in larvae are specified into adult RGC types in a gradual but possibly asynchronous fashion (**Figure 3I**). This late diversification may underlie the post-larval emergence of visual behaviors such as shoaling and visual recognition of places, which are not observed until juvenile stages (Larsch and Baier, 2018; Yashina et al., 2019).

### An intersectional strategy enables genetic access to RGC types

To relate transcriptional clusters to RGC morphology, we chose a set of three TFs with restricted expression to generate transgenic driver lines: *mafaa, tbr1b, and eomesa*. At the larval stage, each TF was expressed in a small number of clusters that formed non-overlapping groups (3 each for *mafaa* and *tbr1b*, 5 for *eomesa*; **Figure 4A**), together encompassing 37% of larval RGCs. Notably, these TFs maintained a robust cluster-specific expression pattern in adults (**Figure 2B**). We used a homology-independent target integration method (Suzuki et al., 2016) (**STAR Methods, Table S3**) to generate driver lines that selectively labeled RGCs expressing *mafaa, tbr1b*, and *eomesa*. In this method, a DNA sequence encoding the transcriptional activator QF2 is knocked into the corresponding TF locus to drive expression of the reporter transgene *Tg(QUAS:loxP-GFP-loxP-epNTR-tagRFP)* hereafter named *Tg(QUAS:switchNTR)*. All of these genes are also expressed outside the retina, often in spatial domains that overlapped with the axonal projection targets (Kunst et al., 2019) (https://fishatlas.neuro.mpg.de/; **Video S1-4**), necessitating an intersectional approach to restrict expression to RGCs (**Figure 4B**). To this end, we intersected each driver line with *Tg(ath5:Cre)* (Förster et al., 2017), wherein Cre recombinase is specifically expressed in RGCs (Kay, 2005) (**Figure S4A-B**), causing TF-expressing RGCs but not other cells to switch from GFP to RFP (**Figure 4C, Figure S4C-E**).

**Figure 4.**
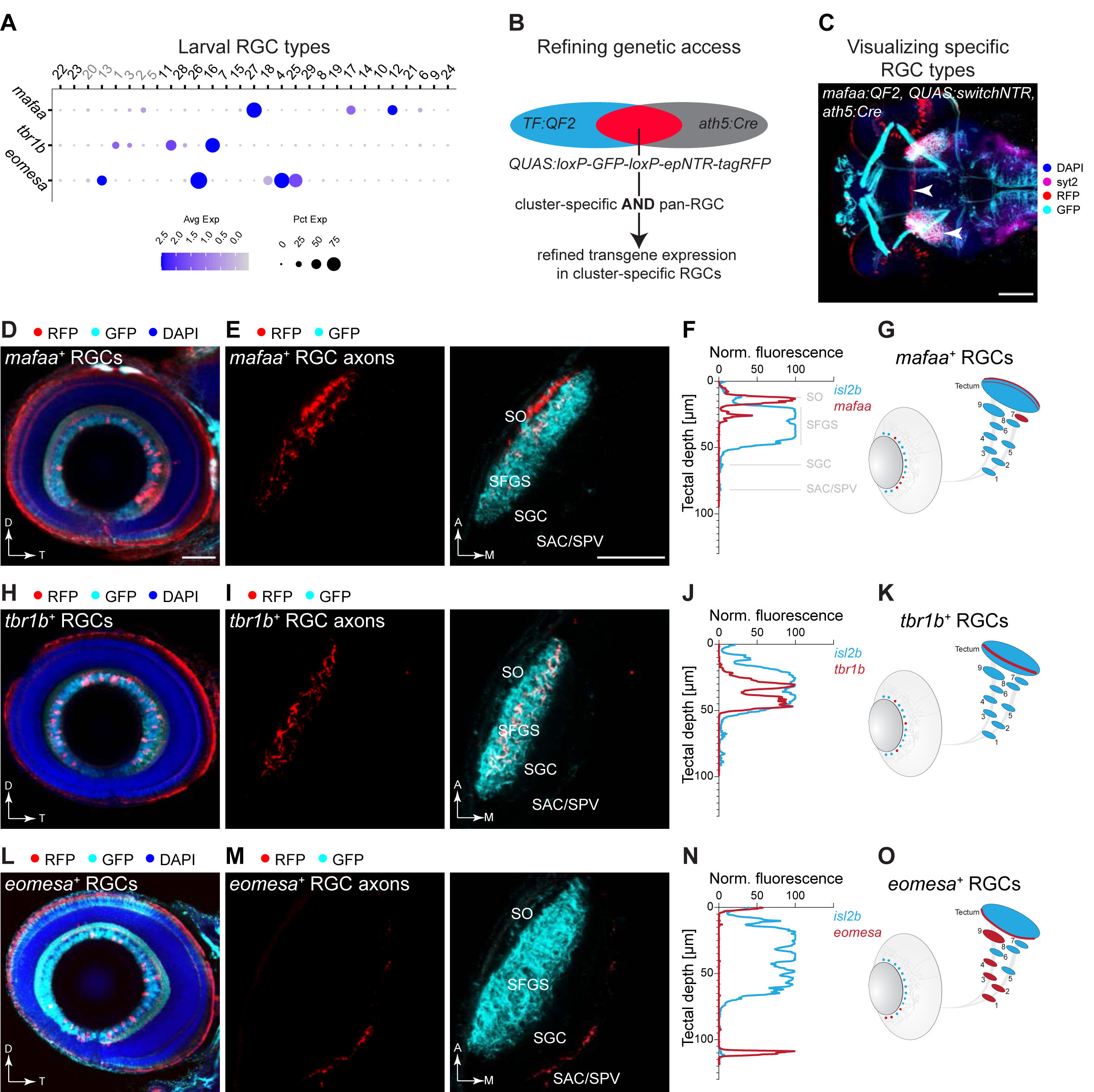
Molecularly defined RGC clusters exhibit distinct axonal projection patterns. **(A)** Dotplot showing expression patterns of *mafaa, tbr1b*, and *eomesa* (rows) in larval clusters (columns) ordered as in **Figure 3A**. Cluster numbers (top) correspond to immature (grey) and mature (black) RGC clusters. **(B)** Marker intersection refines genetic access to TF^*+*^ RGC populations. Briefly, in a driver line TF^*+*^ cells activate expression of GFP through a *QUAS:switchNTR* reporter. Combination with the pan-RGC *Tg(ath5:Cre)* line results in TF^*+*^ RGCs switching to RFP expression, while TF^*+*^ non-RGCs continue to express GFP. **(C)** Immunostained triple-transgenic *Tg(TF:QF2, QUAS:switchNTR, ath5:Cre)* larvae allow for the precise visualization of molecularly defined RGC types as shown here for *mafaa*^*+*^ RGCs (arrows, red labeling). Scale bar, 100 µm. **(D-O)** Anatomical characterization of RGC types labeled by *mafaa* (D-G), *tbr1b* (H-K) and *eomesa* (L-O) using quadruple-transgenic *Tg(TF:QF2, QUAS:switchNTR, ath5:Cre, isl2b:GFP)* larvae. In each case *Tg(isl2b:GFP)* serves as a label for landmarks of RGC projections. Confocal visualization of soma distribution in the immunostained retina (D, H, L), *in vivo* images of axonal projections in the tectum (E, I, M), fluorescence profile across retinotectal laminae measured from the pan-RGC reporter *isl2b* and marker-specific RGC axons (F, J, N) as well as a schematic representation of the soma distribution in the retina and the projection pattern indicating TF^+^ RGCs in red against all RGCs in blue (G, K, O). D, dorsal; T, temporal; A, anterior; M, medial. Scale bar in D for D, H, L, 50 µm. Scale bar in E for E, I, M, 50 µm.

### Molecularly defined RGC subclasses exhibit distinct axonal projection patterns

Previous cell-type classification studies have demonstrated a striking correspondence among the molecular, morphological and physiological properties of retinal neuronal types (Baden et al., 2016; Franke et al., 2017; Peng et al., 2019; Shekhar et al., 2016; Tran et al., 2019; Zeng and Sanes, 2017). In contrast, less is known about the correspondence between these properties of RGC types and the retinorecipient regions to which RGC types project (Dhande et al., 2015; Martersteck et al., 2017). In zebrafish, RGC axons project to ten tectal laminae and nine extratectal arborization fields (AF1-9) (**Figure 1B**). We previously classified zebrafish RGCs at single cell resolution based on stereotyped combinations of dendritic morphologies and axonal projections (Robles et al., 2014). Of note, most (97%) RGCs project axons to the tectum, forming arborizations restricted to a single tectal layer (for nomenclature of the layers, see **Figure 1B**). In addition, about half of the RGCs form *en route* axon collaterals to one or several extratectal targets AF1-AF9. Three percent of RGCs do not reach the tectum, but extend their most distal axon arbor to AF9, the neuropil of the periventricular pretectal nucleus. To ask if axonal patterns of molecularly defined types match individual projection classes described previously (Robles et al., 2014), we crossed the *mafaa, tbr1b* and *eomesa* intersectional lines to *Tg(isl2b:GFP)*, in which all RGCs express GFP. The GFP signal allowed us to compare the intraretinal distribution and axonal projection patterns of RGCs that express each of these three TFs to those that do not.

*mafaa*^*+*^ RGCs were distributed asymmetrically in the retina with a temporal enrichment within the previously described “strike zone”, a zone for high acuity vision, (Bianco et al., 2011; Gahtan, 2005; Mearns et al., 2020; Zhou et al., 2020; Zimmermann et al., 2018) (**Figure 4D**). *mafaa*^*+*^ RGCs showed two projection patterns: They either terminated in the tectal SO, with axon collaterals into AF7, a projection type ascribed to prey-tuned RGCs (Semmelhack et al., 2014); or they arborized exclusively in SFGS2, with no extra AF collaterals, which corresponds to projection class 5 (Robles et al., 2014) (**Figure 4E-G, S4F**). Notably, the innervation of AF7 was heavily biased to the ventral side, suggesting a functional subdivision of this neuropil.

In contrast, *tbr1b*^*+*^ RGCs were distributed throughout the retina (**Figure 4H**). Axons of *tbr1b*^*+*^ RGCs directly innervated intermediate tectal layers (SFGS3 and SFGS5) with no axon collaterals in extra-tectal AFs, corresponding to projection classes 6 and 8, respectively (Robles et al., 2014) (**Figure 4I-K**).

Somata of *eomesa*^*+*^ RGCs were enriched in the ventral retina (**Figure 4L**) and were found to innervate multiple extratectal areas in preoptic area/hypothalamus, thalamus and pretectum (AF1, AF2, AF3, AF4 and AF9) (**Figure S4G**). In the tectum, axons terminated exclusively in the SAC/SPV layer (**Figure 4M-O**), matching projection classes 15 to 20 (Robles et al., 2014). Within the pretectal AF9 neuropil, *eomesa*^*+*^ axon collaterals were restricted to the dorsal half, supporting the previously described subdivision between ventral and dorsal AF9 (Robles et al., 2014).

Together, these results map molecularly defined RGC types to morphologically defined types, and provide a molecular basis for type-specific soma distribution within the retina, as well as projections to retinorecipient nuclei and tectal laminae. Intraretinal asymmetries reflect a functionally biased survey of the visual field, and central projections define the neuroanatomical circuits that process specific visual features.

### RGC types within a subclass exhibit distinct morphologies

We next sought to additionally resolve individual RGC types within the groups defined by *tbr1b* and *eomesa*. The transcription factor *tbx3a* was exclusively expressed by one of the *tbr1b*^*+*^ types (**Figure 5A**). Although the expression level was low in this larval cluster, it was robust and cluster-specific in the corresponding adult cluster (C12; **Figure 1H**). We therefore used the intersectional genome engineering approach to generate a line in which *tbx3a*^*+*^ RGCs were labeled (**Figure S5A**). The line sparsely labeled *tbx3a*^*+*^ RGCs, which demonstrated diffuse dendrites corresponding to the D2 type (Robles et al., 2014) (**Figure 5B**). The morphological features of *tbx3a*^*+*^ RGCs were consistent with them being a subset of *tbr1b*^*+*^ RGCs. *tbx3a*^*+*^ axons terminated exclusively in SFGS5 (**Figures 5C-D**), which is one of the tectal layers to which *tbr1b*^*+*^ RGCs project (**Figures 4I-K**).

**Figure 5.**
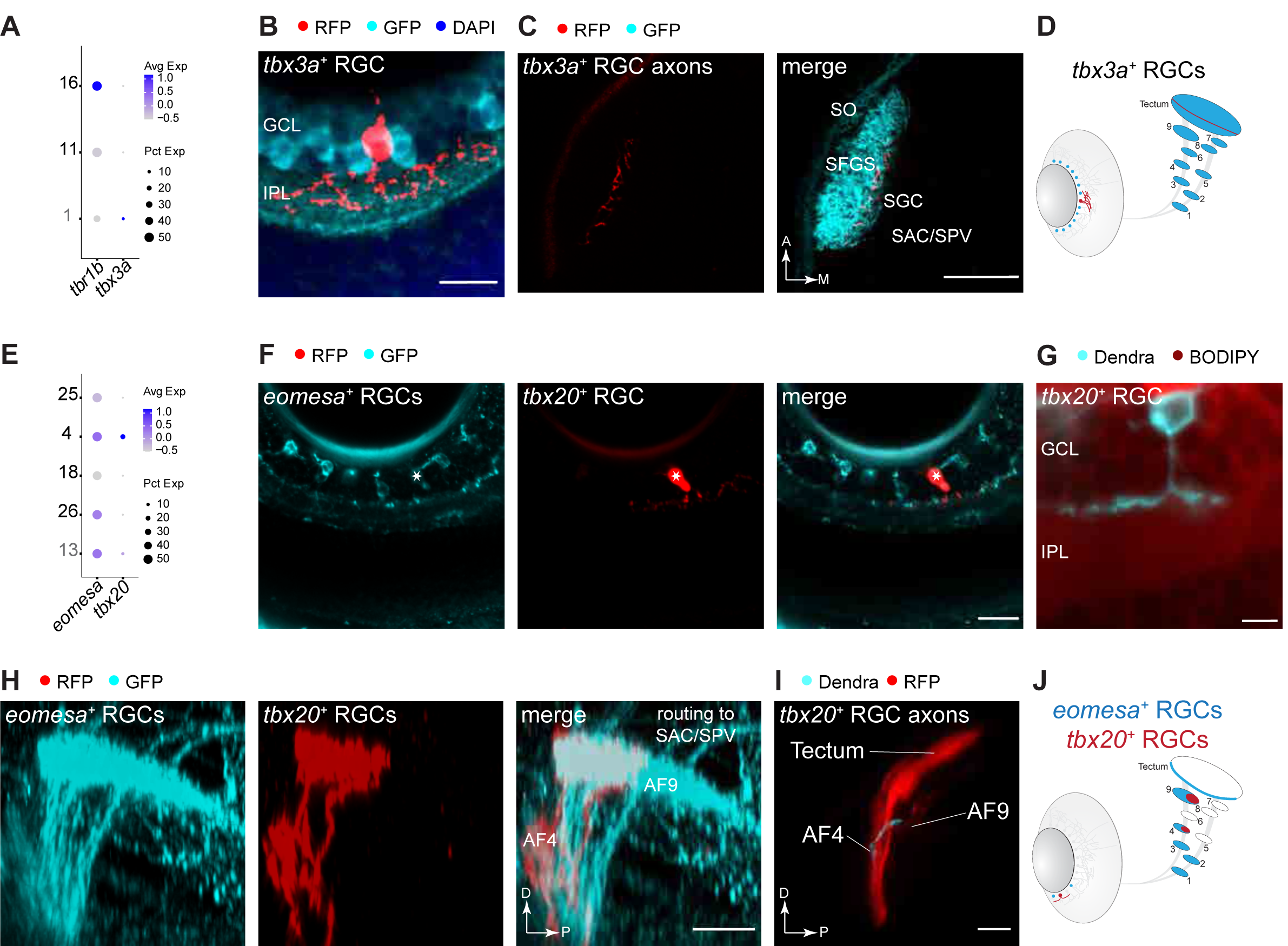
Morphological differences within RGC subclasses. **(A)** Dotplot showing selective co-expression of *tbx3a* in larval *tbr1b*^*+*^ cluster 1. **(B)** Immunostained retina of a *Tg(tbx3a:QF2, QUAS:switchNTR, ath5:Cre, isl2b:GFP)* larva shows diffuse dendrites of *tbx3a*^*+*^ RGCs in the retinal IPL. GCL, ganglion cell layer; IPL, inner plexiform layer. Scale bar, 10 µm. **(C)** *in vivo* confocal plane of *Tg(tbx3a:QF2, QUAS:switchNTR, ath5:Cre, isl2b:GFP)* larvae shows *tbx3a*^*+*^ RGC axons terminating in a deep SFGS layer. A, anterior; M, medial. Scale bar, 50 µm. **(D)** Schematic representation of the soma distribution and projection patterns of the RGC type labeled by *tbx3a* (red) against all RGCs (blue). **(E)** Dotplot showing specific expression of *tbx20* in larval *eomesa*^*+*^ cluster 4. **(F)** Immunostained retinal section of a quadruple-transgenic *Tg(eomesa:QF2, QUAS:GFP, tbx20:Gal4, UAS:NTR-mCherry)* larva showing GFP-labeled *eomesa*^*+*^ RGCs (left), one of which also expresses *tbx20*^*+*^ based on RFP-staining (right, star indicates the co-labeled cell). Scale bar, 20 µm. **(G)** Confocal plane of a live *Tg(tbx20:Gal4, UAS:Dendra)* larval retina with a BODIPY neuropil counterstain. *tbx20*^*+*^ RGCs exhibit monostratified dendrites in the ON IPL. GCL, ganglion cell layer; IPL, inner plexiform layer. Scale bar, 5 µm. **(H)** Pretectal area of a quadruple-transgenic fish with GFP-immunostained *eomesa*^*+*^ RGC axons and RFP-immunostained *eomesa*^*+*^*tbx20*^*+*^ RGC axons, which innervate AF4 and terminate in AF9. D, dorsal; P, posterior. Scale bar, 20 µm. **(I)** 3D side view of the optic tract imaged from a live *Tg(tbx20:Gal4, UAS:Dendra, isl2b:tagRFP)* larva showing that *tbx20*^*+*^ RGC axons innervate AF4 and terminate in AF9. D, dorsal; P, posterior. Scale bar, 50 µm. **(J)** Schematic of soma distribution and axon projections of the RGC type labeled by *tbx20* (red) against all *eomesa*^*+*^ RGCs (blue).

One of the five *eomesa*^*+*^ RGC types specifically expressed the transcription factor *tbx20* (**Figure 5E**). We labeled the *eomesa*^*+*^*tbx20*^*+*^ RGCs by crossing *Tg(eomesa:QF2, QUAS:switchNTR)* to *Tg(tbx20:Gal4, UAS:NTR-mCherry)* (Förster et al., 2017) (**Figure S5B**). The subset of *eomesa*^+^ RGCs that was *tbx20*^*+*^ had monostratified dendrites in the ON layer (**Figure 5F, G**) and axons that extended collaterals into AF4 and terminated in AF9, corresponding to projection class 15 (Robles et al., 2014) (**Figure 5H-J**). In contrast, *eomesa*^*+*^*tbx20*^*-*^ RGCs projected further to the tectal SAC/SPV layer. Together, these results demonstrate that types within a molecularly defined subclass can have distinct dendritic and innervation profiles.

### Molecularly defined RGC types form visual feature-specific channels

To ask whether molecularly distinct RGC types also exhibit distinct functional properties, we harnessed *eomesa* and *tbx20* as markers for an RGC subclass and a unique type within that subclass, respectively. We expressed the cytosolic calcium sensor GCaMP6s in RGCs and recorded axonal calcium transients in response to a set of visual stimuli that included luminance changes (bright and dark flashes), and patterned stimuli such as moving gratings, prey-like and looming stimuli (**STAR Methods, Figure 6A**). Because *tbx20*^*+*^ RGCs do not innervate the tectum, we focused our initial functional analysis on RGC axons in pretectal AF9 (**Figure 6B-D**).

**Figure 6.**
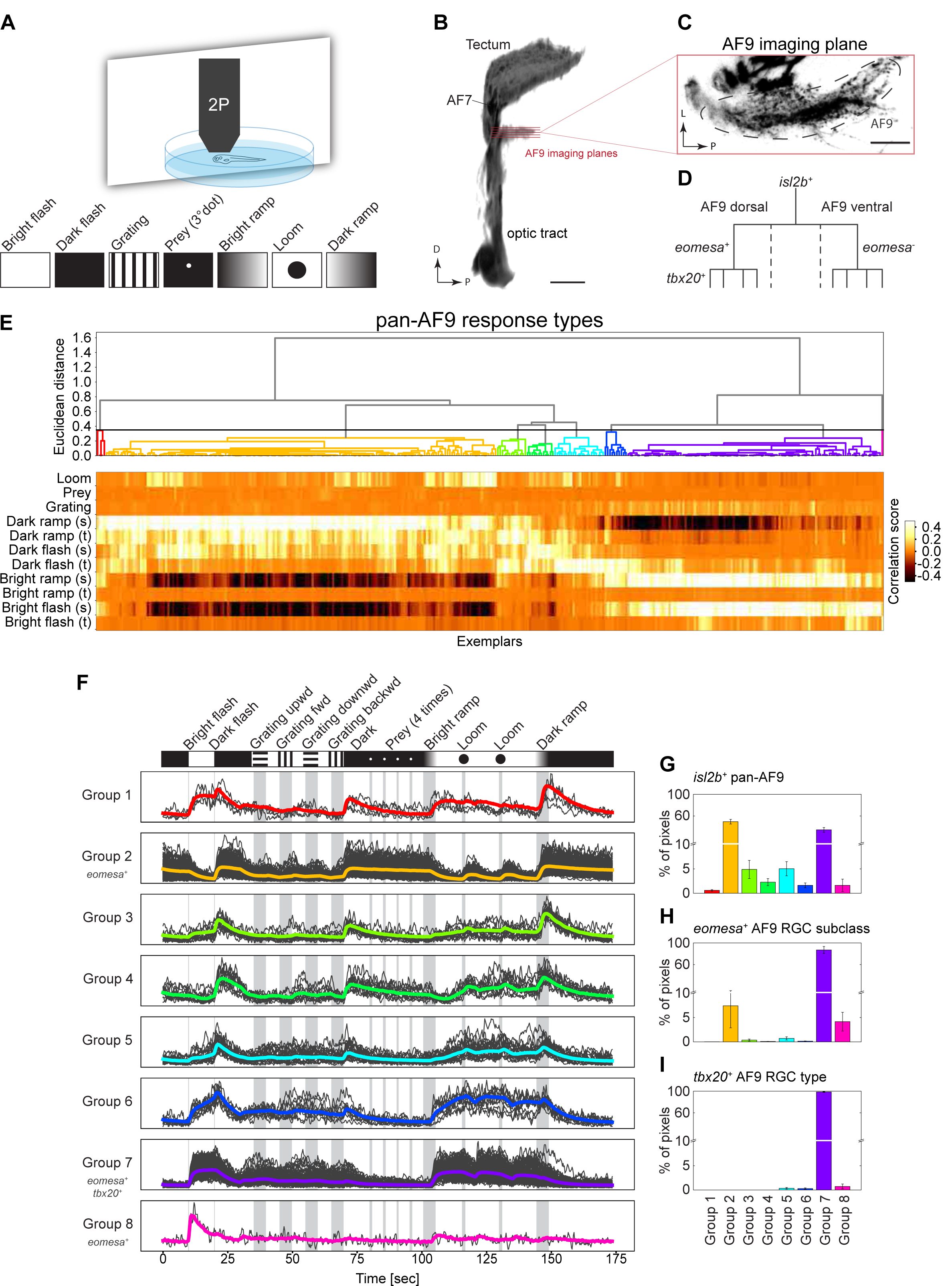
RGC types exhibit specific physiological profiles. **(A)** Experimental setup to characterize tuning profiles of RGC types. Neuronal activity was recorded from immobilized larvae expressing the calcium sensor GCaMP6s in RGC axons during presentation of different visual stimuli using 2P microscopy. **(B)** 3D projection of the optic tract indicating the imaging planes in both ventral and dorsal subdivisions of the pretectal RGC neuropil AF9. D, dorsal; P, posterior. Scale bar, 50 µm. **(C)** Single imaging plane in AF9. L, lateral; P, posterior. Scale bar, 20 µm. **(D)** Hierarchical illustration of genetically defined RGC populations used for functional imaging experiments: *isl2b* labels all RGCs in AF9, *eomesa* marks a subclass in dorsal AF9, wherein *tbx20* is expressed by a unique type among eomesa+ RGCs. **(E)** Characterization of the baseline diversity of *isl2b*^*+*^ RGC responses to visual stimuli as measured in pretectal AF9 axons. Neural activity recordings derived from single pixels were clustered using affinity propagation to reduce noise, resulting in 345 clusters represented by “exemplars” (**STAR Methods**). Hierarchical clustering was used to classify exemplar activity into eight major response groups (dendrogram, top). Heatmap (bottom) depicts activity correlation of exemplars (columns) to each component of visual stimulus (rows). Sustained (s) and transient (t) activity was observed in responses to changing luminance levels. **(F)** Activity traces of eight classified response groups shown in E to the stimulus sequence. Shown are the averaged traces (colored lines) and all representing exemplars that fall into the group (grey lines). Response groups encompass different numbers of exemplars depending on their abundance. **(G-I)** Relative frequencies of the eight response groups in *isl2b*^*+*^ RGCs (G), *eomesa*^*+*^ RGCs (H) and *tbx20*^*+*^ RGCs (I).

We began by characterizing the baseline diversity responses in AF9 by measuring activity from axons in the *Tg(isl2b:Gal4, UAS:GCamP6s)* line, which labels nearly all RGCs (**Figures 1G, 3A**). Regression and clustering analysis classified RGC responses based on their activity to the stimulus components into 8 distinct pretectal response groups (**STAR Methods, Figure 6E**). By visual inspection, we categorized these groups as “sustained ON” (cluster 7), “transient ON” (cluster 8), “sustained OFF” (cluster 2,4) or “sustained ON – transient OFF” (cluster 1,3,5,6). AF9-projecting RGCs were insensitive to patterned stimuli but responded robustly to luminance changes (**Figure 6F**). The relative distribution of these responses varied with sustained ON and OFF types (groups 2 and 7) predominating (**Figure 6G and S6A**).

We next recorded from *eomesa*^*+*^ RGCs. 87% of *eomesa*^*+*^ RGCs mapped to response group 7 (“sustained ON”; **STAR Methods, Figure 6H and S6B**), suggesting that this group of RGCs predominantly encodes ambient luminance levels. The remaining *eomesa*^*+*^ RGCs, mapped to response groups 2 (7%) and 8 (4%). Finally, *eomesa*^*+*^*tbx20*^*+*^ RGCs, mapped almost exclusively to response group 7 (**Figure 6I and S6C**), demonstrating a tight correspondence between molecular, morphological and physiological properties. Thus, the observed functional specification aligns to our molecular definitions, with *tbx20*^*+*^ RGCs representing a unique type of *eomesa*^*+*^ RGCs, which in turn form a subset of *isl2b*^*+*^ RGCs.

To ask whether the diversity of responses in AF9 represent the full range of response types, we recorded from the retinotectal laminae of *Tg(isl2b:Gal4, UAS:GCaMP6s)* larvae. Using the computational pipeline applied to AF9, we observed 14 response groups. The retinotectal RGCs also responded to the presentation of visual objects, such as prey-like stimuli, and their direction of movement, features that are invisible to AF9 (**Figure S6D-F**). Interestingly, tectal laminae received input from distinct response groups, which is attributable to dedicated innervation by unique sets of RGC types (**Figure S6G**). These results demonstrate that specific visual representations are relayed to distinct retinorecipient brain areas, with AF9 receiving a subset of the overall visual information.

### *eomesa*^+^ RGCs express melanopsin

ipRGCs are an evolutionarily ancient, photosensitive subclass of RGCs that were originally implicated as mediators of non-image forming functions such as circadian entrainment and pupillary light reflex, but also play roles in image processing (Do, 2019; Fu et al., 2005; Hattar, 2002; Schmidt et al., 2011). In mammals, *Eomes* (also known as *Tbr2*), the ortholog of *eomesa*, is expressed selectively although not exclusively by ipRGCs (Mao et al., 2014; Peng et al., 2019; Sweeney et al., 2014; Tran et al., 2019), raising the possibility that at least some *eomesa*^*+*^ RGCs in zebrafish are ipRGCs. The canonical marker of ipRGCs is the photopigment melanopsin (*Opn4)* (Berson et al., 2002; Do, 2019; Gooley et al., 2001). We therefore assessed expression of *opn4* in our dataset. While mammals have a single melanopsin gene (*Opn4*), zebrafish contain five melanopsin homologs, three of which (*opn4*.*1, opn4a* and *opn4b*) are related to mammalian *Opn4* and two (*opn4xa* and *opn4xb*) are more closely related to the Xenopus gene *Opn4x* (Bellingham et al., 2006; Matos-Cruz et al., 2011). Previous studies found zebrafish melanopsin genes to be expressed across multiple cell classes in the larval retina, but only *opn4xa* was expressed in RGCs (Matos-Cruz et al., 2011; Zhang et al., 2017). Surprisingly, a large proportion of RNA-seq reads for four of these genes mapped to presumptive intronic locations within the corresponding gene loci (**Figure S7A**). The reads were concentrated in short sequences within the introns, suggesting that they could be derived from unannotated exons (**Figure S7B**) (see Zhang et al., 2020). In fact, in each case, the sequence encoded at least one open reading frame, although the predicted amino acid sequence displayed no significant homology to opsins in any species (data not shown). Nonetheless we combined the intronic and exonic reads to examine cell-type specific expression patterns (La Manno et al., 2018) (**STAR Methods**). This analysis suggested that while *opn4*.*1, opn4a* and *opn4xb* were largely expressed in non-RGCs (**Figure S7C**), *opn4xa* and *opn4b* were robustly enriched in RGCs at both the larval and adult stages. Strikingly, within RGCs, their expression was restricted to *eomesa*^*+*^ RGC types and an additional type (**Figure 7A, S7D**). Taken together, these results provide strong evidence that *eomesa*^*+*^ RGCs types contain ipRGCs.

**Figure 7.**
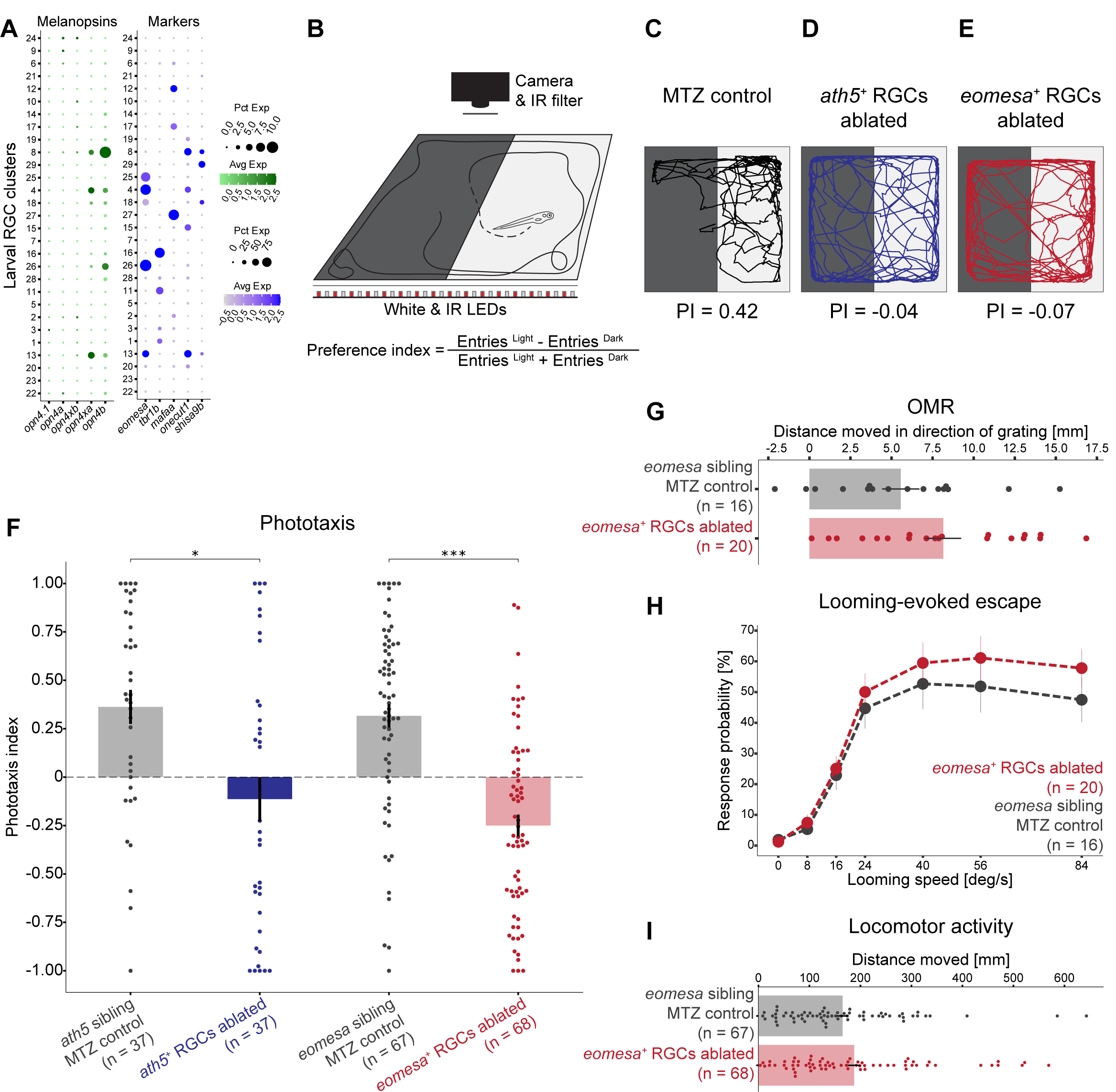
*eomesa*^*+*^ RGCs regulate phototaxis. **(A)** Dotplots showing type-specific expression of melanopsin in larval RGCs. *Left*: Of the five melanopsin homologs (columns), only *opn4xa* and *opn4b* have discernible expression in specific larval RGC clusters (rows). *Right: opn4xa* and *opn4b* expressing clusters include *eomesa*^*+*^ RGC types but not *mafaa*^*+*^ or *tbr1b*^*+*^ types. The only *opn*^+^*eomesa*^*-*^ RGC type is marked by the co-expression of *onecut1* and *shisa9b*. Larval clusters are ordered as in **Figure 3A**. These expression patterns are conserved in adult RGC types (**Figure S7D**). **(B)** Experimental setup used to assay phototactic behavior. Larvae are placed in a light-dark choice arena and their positions are tracked over time. A phototaxis index (PI) is calculated to quantify attraction towards the light source. **(C-E)** Representative traces of a MTZ-treated control larva (C), *ath5*^*+*^ RGC ablated blind larva (D) and *eomesa*^*+*^ RGC ablated larva (E) with indicated PI values. **(F)** PI values for all tested groups: NTR^-^ *Tg(ath5:QF2)* and *Tg(eomesa:QF2)* control siblings as well as *ath5*^*+*^ RGC ablated blind fish and *eomesa*^*+*^ RGC ablated larvae. Each dot represents one fish. Error bars represent SEM. * p<0.05, *** p<0.001 (Dunn post-hoc test). **(G)** Quantification of optomotor response in MTZ-treated control siblings and *eomesa*^*+*^ RGC ablated larvae. Each dot represents one fish. Error bars represent SEM. **(H)** Response probability of MTZ-treated control siblings and *eomesa*^*+*^ RGC ablated larvae to an escape-evoking looming disc. Each dot represents the mean value at a given stimulus expansion rate. Error bars represent SEM. **(I)** Quantification of locomotor activity of MTZ-treated control siblings and *eomesa*^*+*^ RGC ablated larvae. Error bars represent SEM.

### *eomesa*^+^ RGCs are required for phototaxis

To test if *eomesa*^*+*^ RGCs subserve a specific behavior, we selectively ablated them and tested fish in a battery of behavioral assays: (a) phototaxis, the tendency to move towards a light source; (b) the optomotor response (OMR), a reflexive behavior that stabilizes the animal’s position in response to optic flow; (c) escape behavior evoked by a looming stimulus; and (d) overall locomotor activity. In each case, we used *Tg(eomesa:QF2, QUAS:switchNTR, ath5:Cre)* larvae, in which the enzyme nitroreductase (NTR) is specifically expressed by *eomesa*^*+*^ RGCs. NTR converts the substrate metronidazole (MTZ) into a cytotoxic compound (Curado et al., 2008; Tabor et al., 2014). Administration of MTZ to larvae of this strain effectively and selectively ablated *eomesa*^*+*^ RGCs (**STAR Methods, Figure S7E**).

To examine phototaxis, we tracked the position of larvae in a dark-light choice arena and calculated a preference index (PI) (**STAR Methods, Figure 7B**). Control larvae preferred the illuminated half of the arena, quantified as positive phototaxis (**Figures 7C, 7F**; PI 0.36 ± 0.08 for *ath5:QF2* clutch siblings, PI 0.32 ± 0.06 for *eomesa:QF2* clutch siblings, mean ± SEM). As expected, phototaxis was severely disrupted upon NTR activation in a pan-RGC line, a treatment that lesioned all RGCs. (**Figures 7D, 7F**; PI -0.11 ± 0.11, p=0.013, Dunn’s test with Bonferroni correction). Phototaxis was similarly abolished upon selective ablation of *eomesa*^*+*^ RGCs (**Figure 7E, F and S7F**; PI -0.25 ± 0.06, p=1.176 × 10^−7^, Dunn’s test with Bonferroni correction). In contrast, we found no significant effect of removing *eomesa*^*+*^ RGCs on OMR, escape behavior or overall locomotor activity (**STAR Methods, Figure 7G-I**). In conclusion, this defined RGC subclass mediates phototaxis, providing compelling evidence of a visual pathway regulating a specific behavior.

## Discussion

Using single cell transcriptomics, we assembled a comprehensive molecular catalog of RGC types in adult and larval zebrafish. We identified a number of transcription factors (TFs), cell recognition molecules, and neuropeptides that were expressed in one or few of the RGC clusters, serving as single or combinatorial markers for individual cell types. We note that orthologs of many variably expressed TFs in zebrafish also selectively label RGC types in mouse, macaques and human (e.g. *Pou4f3, Eomes, Tbr1, Irx3, Mafa, Satb1, Satb2, Foxp2*) (Liu et al., 2018; Peng et al., 2019, 2017; Rousso et al., 2016; Tran et al., 2019; Yan et al., 2020). This observation highlights the conservation of TF expression patterns and potentially cell type identities across the vertebrate lineage. Based on this parallel, we chose four type-specific TFs to genome-engineer driver lines and used them to investigate the morphological and physiological properties of specific RGC types. Finally, we exploited this precise genetic access to causally associate a molecularly, anatomically and functionally defined retinofugal pathway with a specific visual behavior. Together, our work provides a comprehensive survey of RGC diversity as a resource for future studies and paves the way to holistically explore the development, structure and function of the vertebrate visual system.

### Diversity of zebrafish RGCs

We identified 33 transcriptionally distinct RGC types in adult zebrafish. The number and frequency distribution of cell types bears resemblance to those seen in other vertebrates like mice (∼40-46 types: Rheaume et al., 2018; Tran et al., 2019), Peromyscus and chick (Sanes and co-workers, *unpublished*). On the other hand, the primate and human visual system contains only 15 to 18 molecular RGC types (Peng et al., 2019; Yan et al., 2020) with the four most frequent types, the so called ON and OFF midget and parasol RGCs, accounting for ∼85% of all RGCs. In contrast, the four most frequent RGC types in zebrafish, mice, chick and Peromyscus account for <30% of all RGCs.

Strikingly, the number of molecular types identified in this study (33) is substantially lower than the >50 types described previously based on morphology alone (Robles et al., 2014). In fact, the latter study recorded 75 distinct combinations of dendrite stratification and axonal projection patterns from a collection of 450 individually traced larval RGCs. As retinal projectomes are not available for other vertebrate species, one can only speculate on the reasons for this discrepancy. Our transcriptomic catalog may underestimate the true diversity for the following reasons: (a) Some types may have been undersampled because of biases in expression of the transgene or selective vulnerability in the purification process. (b) Some infrequent types might be unresolved, because the computational power to resolve types into separate clusters depends on transcriptional separation and the number of cells sequenced (Shekhar et al., 2016). (c) Distinctive transcriptional signatures may exist only during very early development and are downregulated at the stages we sampled, as observed in previous studies (Li et al., 2017; Tran et al., 2019). Using more sensitive RNA sequencing methods or measuring other modalities (e.g. proteome or epigenome) might potentially resolve these “hidden” types. On the other hand, the projectome study may have overestimated the cell type diversity, as some morphologies might be developmentally transient and be pruned or error-corrected at later stages. Taken together, it seems plausible that the true number of RGC types in zebrafish is greater than 33 and fewer than 75. Continued efforts to map the transcriptional profiles of specific types to their morphology, function and distribution in the retina, at different stages of development, will settle this issue and produce a definitive account of RGC diversity.

### Progressive diversification of RGCs

Unlike in mammals and birds, the teleost retina grows throughout life, constantly adding new RGCs at its margins, although expansion and neurogenesis rates drop in the adult (Marcus et al., 1999). Against this backdrop of continued growth and rewiring (Hu and Easter, 1999; Kay, 2005), the zebrafish visual system supports a variety of behaviors already at early larval stages. We therefore expected to find a mix of fully differentiated and immature RGCs, especially in our larval dataset. Because most larval behaviors persist into adulthood, we also hypothesized that many larval RGC types would have matching counterparts in the adult. Comparison of the larval and adult RGC clusters confirmed both of these hypotheses. Twenty-three larval clusters, containing two-thirds of the RGCs, could be successfully mapped onto one or very few adult clusters. Attempts to further subdivide multimapping larval clusters were unsuccessful, suggesting that molecular diversification may be incomplete for these types at the larval stage. A third of the larval RGCs, belonging to 6 of the 29 larval clusters, displayed signatures of ongoing differentiation, including genes associated with cytoskeletal reorganization and axon guidance. Five of the immature clusters exhibited transcriptomic signatures that mapped onto distinct subsets of mature larval RGC types, indicating that they are cells in transition to maturity. The sixth cluster may represent postmitotic RGC precursors that are apparently not yet committed to specific fates. These immature RGCs persist into adulthood (adult cluster C4). However, such immature RGCs were not detected in adult mouse, primate or human RGCs (Macosko et al., 2015; Peng et al., 2019; Rheaume et al., 2018; Shekhar et al., 2016; Tran et al., 2019; Yan et al., 2020).

These results suggest a model in which RGC types arise by progressive diversification, proceeding from a precursor, through immature and incompletely specified larval types that finally mature into specific adult types. A corollary is that not all RGC types are completely specified in larvae. Those that do form later may support behaviors that emerge at juvenile stages, such as shoaling (Larsch and Baier, 2018) or spatial navigation based on visual cues (Yashina et al., 2019). This diversification proceeds in parallel with global as well as type-specific gene expression changes even in the stable clusters. Nonetheless, we find potential larval counterparts of all adult types, suggesting a progressive addition of new types without loss of previously formed ones.

### Matched molecular, physiological and morphological properties of RGC types

One of the central goals in classifying the diverse cell types that comprise the nervous system is to harmonize multiple aspects of cell identity (Regev et al., 2017; Sanes and Masland, 2015; Vlasits et al., 2019; Zeng and Sanes, 2017). Recent studies in mice have demonstrated congruence between molecularly, physiologically and morphologically defined RGC types (Baden et al., 2016; Bae et al., 2018; Tran et al., 2019), although none of these studies assayed all properties together. Also, morphological characterization was restricted to dendrites, excluding axonal projections (Bae et al., 2018). Using larval zebrafish as a model system enabled us to survey both structural and functional properties of molecularly defined RGC types. Our results suggest that these features are tightly intertwined.

By exploiting selectively expressed TFs in our catalog, we engineered reporter lines that target either an individual or a small group of closely related RGC types. In each case, we found labeling of an anatomically distinct visual pathway with dedicated projection targets in the brain. *mafaa*^*+*^ RGCs target AF7 and SO, the most superficial tectal layer; *tbr1b*^*+*^ RGCs innervate the mid tectal domain; and *eomesa*^*+*^ RGCs project to SAC/SPV, the deepest layer of the tectum. Intriguingly, RGCs resembling the *mafaa*^*+*^ projection pattern have previously been implicated in the recognition of small, motile prey (Semmelhack et al., 2014). Molecular and morphological characteristics also mapped to discrete physiological tuning profiles. For example, *tbx20*^*+*^ RGCs, which constitute a rare RGC type that projects to two pretectal nuclei (AF4 and AF9), but does not extend a collateral arbor to the tectum, possess transcriptomic, morphological and activity patterns that are consistent with them being a single type within the group of *eomesa*^*+*^ RGCs. We speculate that *eomesa*^*+*^ RGC-specific expression of the secreted morphogen *bmp4* and axon guidance receptor *plxna4* contribute to their asymmetrical position within the retina and their axonal projection patterns within the brain, respectively. In addition, this group of RGCs may exert neuropeptidergic functions via their synthesis of the neuropeptide *nmbb*. These hypotheses can now be tested, using the genetic access provided by our work.

### Specific behavioral role of eomesa^+^ RGCs

For behavioral studies, we focused on *eomesa*^*+*^ RGCs. These RGCs co-expressed not only *opn4xa* as previously suggested (Matos-Cruz et al., 2011), but also *opn4b*, consistent with the idea that they are ipRGCs. *Tbr2*, the mammalian homolog of *eomesa*, is expressed in mouse RGCs and essential for their establishment and maintenance (Mao et al., 2014).

In mammals, melanopsin confers the ability to sense ambient light stimuli (Berson et al., 2002; Do, 2019; Schmidt et al., 2011). Similarly, zebrafish melanopsin-expressing RGCs have previously been linked to phototaxis (Zhang et al., 2017), but lack of precise genetic access has precluded a direct test of this connection. Functional imaging during presentation with a battery of visual stimuli revealed that *eomesa*^*+*^ RGCs specifically encode ambient luminance levels. By selectively ablating *eomesa*^*+*^ RGCs, we showed that they are required for phototaxis but dispensable for several other visually guided behaviors. These results add to the growing consensus that genetically defined and anatomically separable axonal pathways convey specific visual features to downstream processing centers for initiation of appropriate behavioral responses. Such functional specification of visual pathways is also evident in the mouse visual system (Baden et al., 2016; Bae et al., 2018; Chen et al., 2011; Dhande et al., 2013; Freedman, 1999; Güler et al., 2008; Hattar et al., 2003; Piscopo et al., 2013; Sanes and Masland, 2015).

In conclusion, our investigations of the functional pathways carried by individual, molecularly defined RGC types support a ‘labeled line’ architecture of the visual system. Neural circuits downstream of the tectum and the other retinorecipient nuclei in the hypothalamus, thalamus and pretectum may be organized in a similar fashion. For example, the tectal motor map was recently shown to channel commands to hindbrain circuits via at least two parallel pathways, each dedicated to a specific behavioral response (Helmbrecht et al., 2018). Our scRNA-seq approach to RGC diversity is expected to serve as a blueprint for the molecular dissection of other parts of the central nervous system.

## Supporting information

Video_S_eomesa

Video_S_mafaa

Video_S_tbr1b

Video_S_tbx3a

## Acknowledgements

Funding for this study was provided by the Max Planck Society (YK, TOH, MS, AMF, SL, EL, IAA, HB), the DFG-SPP1926 (AMF, HB), NS029169, National Institute of Health grants EY022073 (JRS) and R00EY028625 (KS), and startup funding from UC Berkeley (KS, JH, SB). We would like to acknowledge the Graduate School of Systemic Neurosciences, GSN-LMU Munich, for supporting this project by travel grants. The authors would like to thank all members of the Sanes and Baier labs and colleagues at the Max Planck Institute of Neurobiology for discussions and comments. We are grateful to Johannes Larsch, Greg Marquart, Joe Donovan, Inbal Shainer and Duncan Mearns for critical reading of the manuscript. We further thank Julia Kuhl for creating scientific illustrations, Robert Kasper at the MPIN imaging facility for technical support, and Irene Whitney, Yirong Peng, Emily Martersteck, Mallory Laboulaye and Dustin Herrmann for experimental guidance and assistance.

## Author contributions

YK, KS, JRS, HB conceptualized the study. YK, MS, TOH, AMF, KS, JH performed experiments and investigated data with help of SL, EL, IAA. AS, JH, KS, AMF, TOH developed computational methods and analyzed data. The manuscript was written by YK, JH, KS, JRS, HB and reviewed by all authors. Funding was acquired by KS, JRS and HB.

## Supplementary Information

### Contact for reagent and resource sharing

Further information and requests for resources and reagents should be directed to and will be fulfilled by the Lead Contact.

### Zebrafish

Adult and larval zebrafish were maintained on a 14:10 hour light:dark cycle at 28°C. Embryos were bred in Danieau’s solution (17 mM NaCl, 2 mM KCl, 0.12 mM MgSO4, 1.8 mM Ca(NO3)2, 1.5 mM HEPES). All animal procedures conformed to the institutional guidelines set by the Max Planck Society, with an animal protocol approved by the regional government (Regierung von Oberbayern) as well as by the Harvard University/Faculty of Arts & Sciences Standing Committee on the Use of Animals in Research and Teaching (IACUC). All animals used were anesthetized in a lethal overdose of tricaine (Sigma, CAT# E10521) and rapidly euthanized by immersion in ice water for 10 min. Zebrafish larvae used in this study were between 5 and 7 days post fertilization.

For single cell transcriptomic profiling, 5 dpf larval and 4-6 months old female and male adult *Tg(isl2b:tagRFP)* (Poulain and Chien, 2013) zebrafish were used for retina dissection, tissue dissociation and cell purification.

To establish intersectional transgenic tools, the *Tg(ath5:Cre)* (Förster et al., 2017) transgene was characterized using *Tg(actb2:loxP-eGFP-loxP-lynTagRFPT)* (Marquart et al., 2015) fish.

For morphological analysis of specific RGC types, the pan-RGC transgenes *Tg(isl2b:tagRFP)* or *Tg(isl2b:GFP)* (Pittman et al., 2008) served as a visual landmark of target brain nuclei. We generated the RGC type-specific transgenic lines *Tg(eomesa:QF2), Tg(mafaa:QF2), Tg(tbr1b:QF2), Tg(tbx3a:QF2)* and combined them with *Tg(QUAS:switchNTR)* or *Tg(QUAS:switchG6s)* reporters. In addition, we used *Tg(tbx20:Gal4)* in combination with *Tg(UAS:NTR-mCherry)* (Davison et al., 2007) or *Tg(UAS:Dendra)* (Arrenberg et al., 2009). Larvae were bred in 0.003% PTU (Sigma, CAT# P7629) in Danieau’s to suppress pigmentation prior to staining.

Functional imaging data were obtained from mitfa^-/-^ larvae expressing GCaMP6s in RGCs by crossing *Tg(isl2b:Gal4)* (Fujimoto et al., 2011) or *Tg(tbx20:Gal4)* (Förster et al., 2017) to *Tg(UAS:GCaMP6s)* (Thiele et al., 2014) or generating triple-transgenic *Tg(eomesa:QF2, QUAS:switchG6s, ath5:Cre)* fish.

Targeted cell ablation and subsequent behavioral experiments were performed using *Tg(ath5:QF2, QUAS:epNTR-tagRFP)* (Fernandes et al., 2019) and triple-transgenic *Tg(eomesa:QF2, QUAS:switchNTR, ath5:Cre)* larvae.

## METHODS

### RGC purification and droplet based single cell RNA sequencing

RGCs were labeled using transgenic *Tg(isl2b:tagRFP)* zebrafish that express RFP in all RGCs (Mumm et al., 2006; Pittman et al., 2008). Retinas from larval or adult fish were dissected in oxygenated (ox) Ames (Sigma, CAT# A1420) and transferred into ox Ames on ice until tissue collection was completed. Retinas were digested in papain 20U/ml (Worthington, CAT# LS003126), DNAseI 80U/ml (Sigma, CAT# D4527), L-cysteine 1.5mM (Sigma, CAT# C1276) in ox Ames at 28°C for 30 (larval retinas) or 45 minutes (adult retinas). To stop the digestion, the papain solution was replaced by papain inhibitor solution containing ovomucoid 15mg/ml (Worthington, CAT# LS003087) and BSA (Sigma, CAT# A9418) 15mg/ml. Tissue was gently dissociated by trituration using a flamed glass pipette in papain inhibitor solution. To wash the cell suspension, cells were pelleted at 250g for 8 minutes and resuspended in ox. Ames containing 0.4% BSA. The cell suspension was filtered through a 30µm strainer prior to fluorescence-activated cell sorting (FACS) purification. Non-transgenic wildtype retinas were used to determine background fluorescence levels and adjust sorting gates. Calcein blue (ThermoFisher, CAT# C1429) was added to distinguish live RFP^+^ RGCs. Cells were washed and resuspended in PBS (Gibco, CAT# 10010001) 0.04% BSA and loaded onto the microfluidic device within ∼45 minutes after FACS enrichment. Droplet RNA sequencing experiments using the 10X Genomics chromium platform (Chromium Single Cell 30 Library & Gel Bead Kit v2, CAT# 120237; Chromium Single Cell A Chip Kit, CAT# 1000009; Chromium i7 Multiplex Kit, CAT# 120262) were performed according to the manufacturer’s instructions with no modifications.

For the larval dataset, 200 manually dissected retinas were dissociated in one experiment and single cell profiles were collected across 3 replicates. For the adult dataset, about 20 retinas per batch were dissected and dissociated and droplet RNA sequencing was performed collecting a total of 15 replicates across 5 experiments. The cDNA libraries were sequenced on the Illumina HiSeq 2500 to a depth of ∼30,000 reads per cell.

### Computational analysis of single cell transcriptomics data

#### Alignment and quantification of gene expression

Initial preprocessing was performed using the cellranger software suite (version 2.1.0, 10X Genomics), following steps described previously (Pandey et al., 2018). Briefly, sequencing reads were demultiplexed using “cellranger mkfastq” to obtain a separate set of fastq.gz files for each of 15 adult and 3 larval samples. Reads for each channel were aligned to the zebrafish reference transcriptome (ENSEMBL zv10, release 82) using “cellranger count” with default parameters to obtain a digital gene expression (DGE) matrix (genes x cells) summarizing transcript counts. For each of the adult and larval experiments, we combined the DGEs from different channels and analyzed them, as described below using the Seurat R package (Satija et al., 2015).

#### Adult RGC catalog

##### Preprocessing and batch integration

The adult DGE matrix was filtered to remove genes expressed in fewer than 25 cells, and cells expressing fewer than 450 genes resulting in 24,105 genes and 48,551 cells. To align the five biological replicates, we used the Canonical Correlation Analysis (CCA) based integration framework in Seurat to embed the cells in a shared, reduced dimensional gene expression space. Briefly, each cell was normalized to a total library size of 10,000 and the normalized counts were log-transformed (*X← log*(*X*+ 1)) using the function *Seurat::NormalizeData*. We used *Seurat::FindVariableFetures* with option *selection*.*method = “vst”* to identify the top 2000 highly variable genes (HVGs) (Hafemeister and Satija, 2020) in each batch. Next, we used *Seurat::FindIntegrationAnchors* and *Seurat::IntegrateData*, both with options “dims=1:40” to perform Canonical Correlation Analysis (CCA)-based batch correction on the reduced expression matrix consisting of the HVGs. The “integrated” expression values were combined across batches, and used for dimensionality reduction and clustering.

##### Dimensionality Reduction, Clustering and Visualization

To remove scale disparities between genes arising from differences in average expression levels, the integrated expression values for each HVG were z-scored across the cells using *Seurat::ScaleData*. Next, we performed Principal Component Analysis (PCA) on the scaled matrix, and used *Seurat::ElbowPlot* to select 40 principal components (PCs). In this reduced dimensional space of 40 PCs, we built a k-nearest neighbor graph using *Seurat::FindNeighbors* and identified transcriptionally distinct clusters using *Seurat::FindClusters*, running the Louvain algorithm. Using the top 40 PCs, we also embedded the cells onto a 2D map using t-distributed stochastic neighbor embedding (tSNE) (Linderman et al., 2019; Maaten and Hinton, 2008). These embeddings were used downstream to visualize gene expression patterns as well as the distribution of various metadata (batch ID, cluster ID, cell quality, etc.).

##### Identification of RGCs and filtering contaminant classes

RGC clusters were identified based on expression of the pan-RGC markers *rbpms2b* (Hörnberg et al., 2013) and *isl2b* (Pittman et al., 2008). Clusters were removed if they contained an abnormally low number of average genes per cell, did not express *rbpms2b*, or expressed genes present in contaminant cell types. Examples of such genes include *rlbp1a* and *apoeb* for Muller glia (Bernardos et al., 2007), *vsx1* for bipolar cells (Vitorino et al., 2009), *gad1* and *gad2* for amacrine cells (Sandell et al., 1994), *pde6* for photoreceptors (Abalo et al., 2020), and *cldn19* for endothelial cells (Kolosov et al., 2013). A total of 15,909 cells corresponding to these cell classes were removed. This contamination most likely arises from transgenic labeling of other retinal cells by *Tg(isl2b:tagRFP)* that fall into the same FACS gate as RGCs. Interestingly, we found a much lower proportion of contaminants at the larval stage (see below), which suggests that promiscuous expression of the transgene may increase with age. The RGCs were separated and analyzed beginning from raw counts, with integration, PCA, clustering, and visualization performed in the same way detailed above.

##### Differential expression analysis and hierarchical clustering

We used *Seurat::FindMarkers* with options *test*.*use =“MAST”, max*.*cells*.*per*.*ident = 1000* to identify differentially expressed genes in each RGC cluster. To identify transcriptional relationships between RGC clusters, we used *Seurat::FindVariableFeatures* to recalculate the top 500 most variable genes. The average expression values of genes in each cluster were used as input for hierarchical clustering, performed using *Seurat::BuildClusterTree*. The resulting output was visualized as a dendrogram.

#### Larval RGC catalog

The larval DGE was analyzed by following the steps entitled “*Preprocessing”, “Dimensionality Reduction”, “Clustering”* and “*2D Visualization”* described above. Filtering genes expressed in fewer than 25 cells, and cells expressing fewer than 450 genes resulted in 24,105 genes in 12,698 cells. Each cell was normalized and log-transformed, and the top 2000 HVGs were identified in each batch as before. z-scored expression values along these HVGs were used to calculate PCs, and the top 30 PCs were used to define clusters as well as embed cells on a tSNE map.

We annotated clusters based on their expression of cell-class specific markers as before, and removed non-RGC clusters, which comprised ∼9.3% of the data. We performed differential expression analysis among the RGC clusters (all of which robustly expressed *rbpms2b* and *isl2b*) to define cluster-specific markers. The transcriptional interrelatedness between the larval RGC clusters was visualized on a dendrogram built using hierarchical clustering. Six clusters (1, 2, 3, 5, 13, 20) were identified as “immature” based on shared expression of *alcamb, tmsb*, and *tubb5*, while the remaining 23 clusters were labeled as “mature”. Immature and mature larval RGCs were separately visualized on tSNE maps. To compare the clustering quality between the two subsets, we computed the silhouette score for each cell within each subset. For each point, the silhouette score is defined as *a*(*i*) *b* (*i*) / *max*{*a*(*i*),*b* (*i*)}, where *a*(*i*) is the mean distance between point *i* and all other points in the cluster containing *i* and *b*(*i*) is the minimum mean distance of point *i* to all points in any cluster not containing *i*. Using the *silhouette* function of the R package *cluster* with the tSNE embeddings and cluster labels as inputs, the median silhouette score was computed for each subset.

#### Surveying expression of cluster-enriched Transcription Factors, *Neuropeptides, and Recognition Molecules*

Initial databases of 1,524 transcription factors, 158 neuropeptides, and 387 candidates involved in axon guidance were assembled from the Zebrafish Information Network website (zfin.org) by selecting genes with search terms “transcription factor”, “neuropeptide” or “axon guidance”. The recognition molecule library was expanded from the axon guidance list to 515 genes by searching the larval and adult DGEs for genes that began with *cntn* (Contactins), *eph* (ephrin receptors), *efn* (Ephrin proteins), *robo* (Roundabout family of guidance molecules), *slit* (Slit guidance ligands and receptors), *sema* (Semaphorins), *plx* (Plexins), *nrp* (Neuropilins), *cdh* (Cadherins), *pcdh* (Protocadherins), *ncam* (Neuronal cell adhesion molecules), *cadm* (Cell adhesion molecule genes), and *lrrtm* (Leucine-rich repeat transmembrane proteins). These databases were filtered to include only genes expressed in >30% of cells in at least one cluster within the catalog (adult and larval).

From the database of 1,524 transcription factors, 184 and 186 transcription factors were expressed in the larval and adult RGC catalog, respectively. Of these, 147 candidates were expressed in both the larval and adult datasets, which represents a highly significant overlap unlikely to occur by chance (p < 10^−132^, hypergeometric test). Out of 158 candidate neuropeptides, 10 were expressed in the larva and 11 were expressed in the adult (N=8 shared). Among 515 candidate cell surface and secreted molecules, 67 and 76 were expressed in the larva and adult respectively (N=57 shared). These overlaps were also highly significant based on the hypergeometric test (p < 10^−13^ and p < 10^−44^ respectively).

#### Supervised classification analysis of transcriptional correspondence between catalog and larval RGC types

##### Feature Selection

We first assembled a list of cluster specific markers in both larval and adult RGCs as features for training a multi-class classifier. We applied *Seurat::FindAllMarkers* with arguments “only.pos = TRUE, test.use = ‘wilcox’, min.pct = 0.25, logfc.threshold = 0.25” separately to the adult and larval catalogs to identify genes differentially expressed (DE) within each adult or larval cluster. 641 genes expressed with a significance level of p<10^−10^ in at least one adult or larval cluster were selected as features.

##### Assigning adult identities to larval RGCs

Using the 641 DE genes as features, we trained gradient boosted decision trees on the 23 mature larval clusters. This was implemented using the R package xgboost (Chen and Guestrin, 2016), as in our previous publications (Peng et al., 2019; Tran et al., 2019). For training, expression values along each feature were z-scored to remove scale disparities. We split the larval mature RGCs 60%/40% into training and validation sets, respectively. To avoid overrepresentation of the largest clusters, we capped the representation of each cluster to a maximum of 400 cells. We used the “held-out” labels of the validation set to assess the performance of the classifier, which was found to have an average error rate of 9% (min 0.9%, max 17%) for each of the 23 clusters.

The larval RGC-trained classifier was used to assign a larval identity to each adult RGC based on its expression values along the 641 features. For consistency, the adult RGC expression matrix was z-scored along each of these features. The results were summarized as a confusion matrix, which plots the relative proportion of cells in each adult cluster (rows) that map to each larval cluster (column). Importantly, we note that information regarding the adult clustering was not used in either the training or classification steps.

##### Assigning mature larval identities to immature RGCs

We followed a procedure analogous to the one outlined above, with the exception that the classifier was trained and validated on the 6 immature clusters, and applied to each mature cell.

#### Maturational Changes

A mapping was considered 1:1 if more than 60% of the cells within an adult cluster mapped to a single larval cluster, and that larval cluster received no more than 25% of its mappings from any other adult cluster. Six 1:1 mappings were found in the model. *Seurat::FindMarkers* was used to determine differentially expressed genes between all six adult and larval clusters. A gene was considered to be associated with global maturational changes if the magnitude of its average log fold change (logFC) was greater than 1 and expressed in at least 50% of all larval or adult cells. *Seurat:FindMarkers* was then used to determine differentially expressed genes between each larval and adult cluster that mapped to each other. A gene was considered to be associated with type specific maturational changes if the magnitude of its average logFC was greater than 1, the gene was expressed in at least 50% of either the larval or adult cells, and it was not associated with global maturational changes.

#### Combining intronic and exonic reads to elucidate RGC type-specific expression of melanopsin genes

To include intron aligned reads in the quantification of gene expression, we employed velocyto (La Manno et al., 2018), which uses cellranger-generated binary alignment map (BAM) files to calculate separate DGE matrices corresponding to “spliced” and “unspliced” transcripts by distinguishing between intron-aligned and exon-aligned reads. We ran velocyto on each of the adult and larval samples’ BAM files individually using the following general command line invocation,

~~~
velocyto run -b barcodes.tsv -o /output/path -m mask.gtf bamfile.bam
genes.gtf
~~~

The transcriptome annotation (genes.gtf) file used here was also used for alignment by cellranger. “barcodes.tsv” refers to the list of valid cell barcodes in the sample, an output from the cellranger pipeline. The masking file for suppressing alignment to repetitive elements was downloaded from the UCSC Table Browser (https://genome.ucsc.edu/cgi-bin/hgTables) Sep. 2014 (GRCz10/danRer10) assembly. Velocyto output loom files were processed using in house scripts to compute the spliced and unspliced DGE matrices (DGE_spliced_ and DGE_unspliced_), which were summed into a consolidated expression matrix DGE_tot_ = DGE_spliced_+DGE_unspliced_. We used DGE_tot_ for examining the expression patterns of melanopsin genes in RGC types.

### Establishment of Q-system intersectional transgenic tools

To generate intersectional QUAS plasmids, a Tol2-QUAS; cmlc2:mCherry plasmid (Fernandes et al., 2019) was linearized to insert effector genes. For the QUAS:switchNTR construct, a loxP-GFPcaax-loxP fragment (Förster et al., 2017) and an epNTR-tagRFP fragment (Tabor et al., 2014) were inserted by In-Fusion cloning (Takara, Cat# 638909). Similarly, for the QUAS:switchG6s construct, a loxP-tdTomatocaax-loxP fragment (Förster et al., 2017) and a GCaMP6s fragment (Thiele et al., 2014) were inserted. QUAS reporter lines were generated by Tol2-transgenesis as described previously (Suster et al., 2011).

### Locus-specific transgenesis using CRISPR-Cas9

gRNA target sequences were selected using the CCTop tool (Stemmer et al., 2015). Donor plasmids were cloned using the GoldenGATEway strategy (Kirchmaier et al., 2013) recombining entry vectors carrying fragments with sequences for GBait (pGGEV_-1), target gRNA (pGGEV_2 was mimicked by annealed oligonucleotides), basal promoter e1b (pGGEV_3), QF2-polyA (pGGEV_4), and polyA (pGGEV_5’) into a pGGDestSC_-ATG vector.

CRISPR-Cas9 RNP complex was prepared at a concentration of 1.5 µM as described before (Essner, 2016). Briefly, gRNA was produced by annealing customized crRNA (IDT, Alt-R® CRISPR-Cas9 crRNA) with tracrRNA (IDT, CAT# 1072533) in buffer (IDT, CAT# 11-05-01-12). gRNA was incubated with Cas9 protein (IDT, CAT# 1081060) for 15 minutes at 37°C and donor plasmid was added to a final concentration of 25 ng/µl. The freshly prepared CRISPR-Cas9 cocktail was injected into the cell of transgenic *Tg(QUAS:switchNTR, ath5:Cre)* zygotes. Transient expressors were raised and screened for germline transmission.

### Histological methods

Immunohistochemical staining on whole fish larvae or dissected adult brains was performed following PACT tissue clearing as described previously (Kunst et al., 2019; Treweek et al., 2015). In brief, larvae were fixed in PACT hydrogel monomer solution, deoxygenated and polymerized. Samples were cleared for several days and washed in PBT prior to staining. For adult brains, fixation and clearing time was adapted to 48 hours and two weeks, respectively. After clearing, samples were permeabilized and blocked. Primary antibody incubation (anti-GFP, Invitrogen, CAT# A10262; anti-tagRFP, Invitrogen, CAT# R10367; anti-syt2, ZIRC, AB_10013783) on larval fish took place for 7 days while incubation for adult brains was prolonged to 14 days. Following thorough washing, Alexa-conjugated secondary antibodies and DAPI (Invitrogen, CAT# D1306) were incubated for 3 and 7 days for larval and adult brain samples, respectively. Samples were washed, post-fixed in paraformaldehyde (Alfa Aesar, CAT# 43368) and stored in 87% glycerol. Imaging was performed at a Leica SP8 confocal microscope. Prior to image acquisition, samples were mounted in 2% low melting point agarose in 87% glycerol on bridge slides and coverslipped.

To better characterize labeling in the retina, retinal tissue was sectioned on a cryostat. Transgenic larvae or dissected adult eyes were fixed in 4% PFA in PBT at 4°C overnight, washed and incubated in 35% sucrose in PBST for cryoprotection. Tissue was embedded in TissueTek (Sakura, CAT# 4583), sectioned at 30µm, washed in PBT and blocked in 5% goat serum, 1% BSA, and 1% DMSO in PBT. Staining occurred by incubation with primary antibody for 4 days and secondary antibody for 2 days. Following washing and post-fixation, sections were coverslipped for imaging. Images were processed using the FIJI software (Schindelin et al., 2012).

### Functional imaging and computational methods for characterization of RGC responses

Calcium imaging was performed using transgenic zebrafish larvae expressing GCaMP6s in RGCs on a 2P microscope (Femtonics 3DRC, Femtonics, Tuzlo, Hungary). Larvae were immobilized in 2% low melting point agarose (Invitrogen, CAT# 10143954). The stimulus was projected onto a white diffusive screen (10cm wide, 6cm high) using the red channel of a LED projector. The projection was presented monocularly and covered 120° of the larva’s field of view. Visual stimulation was designed using PsychoPy2 (Peirce et al., 2019) as follows: dark screen (10 sec), bright flash (10 sec), dark flash (10 sec), grating moving in four main cardinal directions (10° black bars interspaced at 30°, 5 sec stationary then 5 sec moving at 20°/sec), dark screen (10 sec), prey-like stimulus (4 repetitions of a 3° bright dot sweeping across the black screen at 90°/sec, 20 seconds), bright ramp (dark to bright red in 5 seconds), bright screen (10 seconds), loom (2 repetitions of a black disc expanding at 30°/second on a white screen, 30 seconds), dark ramp (bright red to dark in 5 seconds), dark screen (10 sec). The total stimulus duration was 160 seconds. Recordings were taken using a 20X 1.0 objective and the laser was driven by a Ti:Sapphire source (Chameleon Ultra II, Coherent) at 920 nm.

Recorded imaging data were pre-processed as described previously (Helmbrecht et al., 2018). In brief, images were motion-corrected using the CaImAn package, uniform filtered over 3 frames and the dF/F was calculated using the 5^th^ percentile of the traces. In total 11 regressors for all stimulus components were created and convolved with a GCaMP6s kernel. Neuronal activity was analyzed pixel-wise by calculating a score of all regressors to the calcium responses of each pixel using a linear regression model of the selected response window with the regressor (Python scikit-learn). For the score, the coefficient of the regression (corresponding to the dF/F) was multiplied by the R2.

To determine overall response types, the scores were normalized per fish to the 99^th^ percentile of all pixels recorded. For the functional clustering of the responsive pixels, 2 *Tg(isl2b:Gal4; UAS:GCaMP6s)* fish (6 planes each, 380 × 178 pixels) were analyzed by first removing pixels with maximum scores smaller than 0.3 (49823 pix). Next, to reduce noise, affinity propagation clustering (scikit learn – preference: median of similarities) was performed. Keeping clusters with at least 50 pixels (0.1 percent of all pixels) yielded in total 345 clusters with chosen exemplars. To extract the global cluster structure, these 345 exemplars were further clustered using hierarchical clustering (scipy.cluster) using correlation as distance metric. Setting a distance threshold of 0.35 classified a total of 42,444 pixels into 8 distinct clusters, each comprising more than 100 pixels. Classification of response types in the tectum presented in Figure S6 was performed similarly and yielded 14 final clusters.

To map response types of genetically-defined RGC populations, pixels from eomesa and tbx20 recordings from five fish each, were first analyzed to calculate the scores to each regressor as described above. Likewise, pixels with maximum scores smaller than 0.3 were removed. A k-nearest neighbor classifier was trained on the isl2b clustered pixel (42,444 pix with cluster labels, k = 100) and the scores of every mapped fish were assigned to the clusters of the isl2b dataset using either predicted labels for the pixel distribution or probability estimates for the population distributions.

### Cell ablation

Larvae expressing the enhanced potency nitroreductase enzyme in RGCs as judged by RFP presence in RGC axons were sorted at 3 dpf. Cell ablation was induced at 4 dpf by bathing larvae in 7.5mM metronidazole (Sigma, CAT# M1547) for 24 hours, followed by continued treatment in 5mM metronidazole for 12 hours. Healthy, well developed larvae showing normal body posture and locomotive activity were then given a recovery period for a minimum of 24 hours and behavioral experiments were carried out at 7 dpf. Successful ablation was confirmed by confocal imaging before and after treatment with metronidazole using randomly selected clutch mates.

### Phototaxis assay

All tests were performed between 9AM and 5PM. Light-preference behavior was assayed by methods modified from (Zhang et al., 2017). Larvae were tested, six animals at a time, in custom-made square chambers (3 × 3 cm each chamber), which was placed inside the ZebraBox (Viewpoint), a device for automated observation and tracking of zebrafish behavior. White light and infrared light were projected from the bottom. Half of a chamber was covered with two dichroic Film Polarizers stacked on top of each other creating a gradient of light intensity near the boundary. The lux intensity of the dark and the light side was around 50 lux and 180 lux, respectively. Animals were allowed to adapt to the arena and light conditions for at least 30 minutes prior to behavioral testing. Calculation of the distance travelled and number of entries to each field was performed by the ZebraBox software. Independent validation of tracking of animals was performed with the TheRealFishTracker (http://www.dgp.toronto.edu/~mccrae/projects/FishTracker/) and results were plotted with custom-written python scripts (Python 3.7) for examples shown on Figure 7. The duration Preference Index was calculated as the difference between the duration (every 10 seconds) in the lit side and the dim side divided by the total duration of the experiment (30 minutes per recording session). The entries PI was calculated as the difference between the number of counts in the lit side and the dim side divided by the total number of counts. Positive PI values indicate light preference. Plots and statistical analysis were performed with custom-written python scripts.

### Looming-evoked escape and optomotor response assay

We adapted previously established methods (Fernandes et al., 2019; Larsch and Baier, 2018) to test optomotor and escape responses. In brief, larvae are placed in glass dishes of 10 cm diameter separated by walls to prevent visual contact. Nine animals could be tested at a time. Diffuse Infrared illumination was provided from below to record animal behavior at 30 fps using a camera equipped with an IR band-pass filter. Visual stimuli were projected onto the projection film from underneath via a cold mirror. Image processing and stimulus generation were performed with Bonsai (Lopes et al., 2015). Before behavioral tests, animals were kept for at least 30 minutes in a petri dish floating above a fully lit portion of the projection screen to allow habituation to light and temperature conditions of the experiment.

Looming stimuli were presented as stationary dots expanding for half a second (15 frames) with a linear increase in diameter. Stimuli were presented 1 cm from the fish at angles of 90° or 270° relative to the animals’ center of mass and orientation at the beginning of the stimulus. Looming stimuli were presented once every 90 seconds. The order of different stimuli (looming expansion rates used: 8, 16, 24, 40, 56 and 84 deg/s, either left or right of the animal) was randomized for each group of animals. To drive larvae towards the center of the dishes using their optomotor response (OMR), a moving grating was presented for 20 seconds ending 10 seconds before the presentation of the next looming stimulus. Distance travelled towards the center during grating presentation was used to measure performance of OMR.

### Data and software availability

Computational scripts detailing scRNA-seq analysis reported in this paper are available at https://github.com/shekharlab/ZebrafishRGC. We have also provided R markdown (Rmd) files that show step-by-step reproduction of the key results. All raw and processed scRNA-seq datasets reported in this study will be released in NCBI’s Gene Expression Omnibus (GEO) under the accession number GSE152842. Visualization of the zebrafish RGC atlas will be made available at https://portals.broadinstitute.org/single_cell (Study ID: SCP992). All data and custom software for functional imaging analysis and behavioral tests will be made available upon request.

## Supplemental Information titles and legends

**Figure S1.**
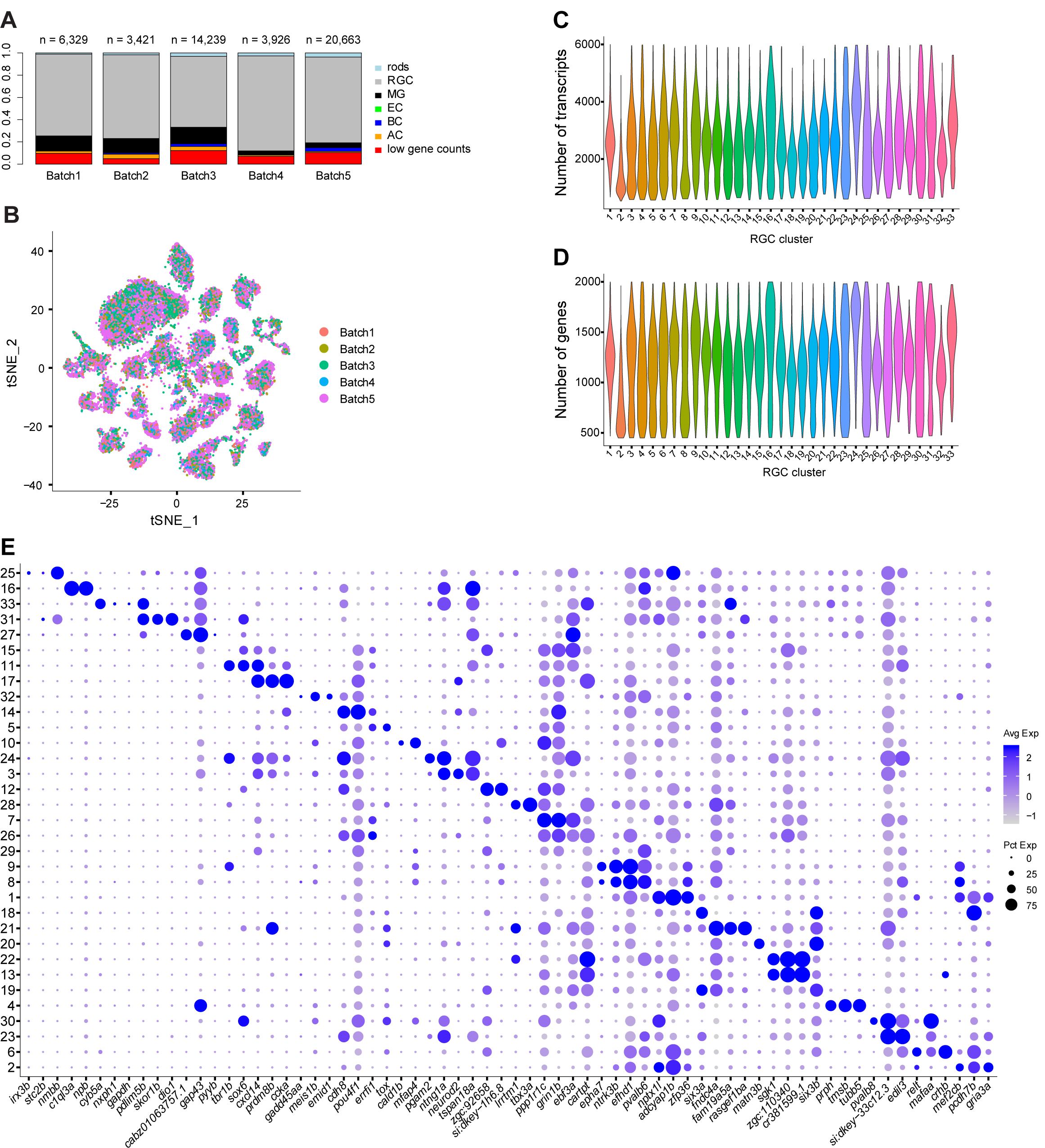
Molecular catalog of adult RGCs, related to Figure 1. **(A)** Barplot showing relative proportions (y-axis) of RGCs, non-RGCs and low-quality cells (colors) in scRNA-seq biological replicates from adult fish (x-axis). Recovered cell numbers from each replicate are indicated on top. non-RGCs include rods, bipolar cells (BC), amacrine cells (AC), Muller glia (MG), and endothelial cells (EC). Clusters containing “cells” with low gene counts that did not exhibit any distinguishing markers were excluded from the analysis. **(B)** tSNE visualization of adult RGCs (x-y coordinates as in **Figure 1E**) colored by replicate ID, showing lack of strong batch effects. **(C)** Violin plot showing distributions of the number of transcripts detected per cell (y-axis) in each adult RGC cluster (x-axis). **(D)** Violin plot showing distributions of the number of genes detected per cell (y-axis) in each adult RGC cluster (x-axis). **(E)** Dotplot showing top three selectively enriched markers (columns) in each adult RGC cluster (rows). Representation and row ordering as in **Figure 1F**.

**Figure S2.**
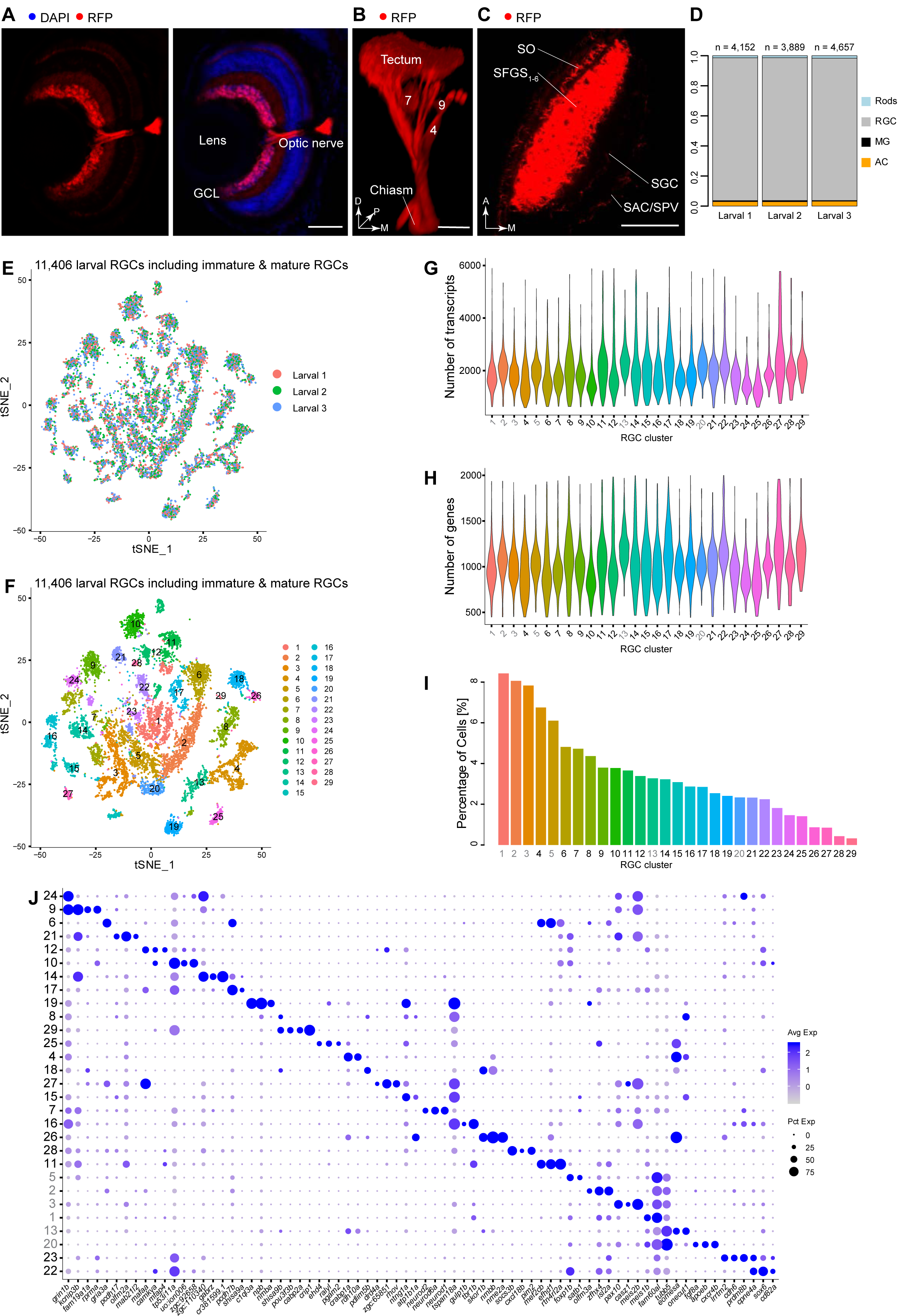
Molecular catalog of larval RGCs, related to Figure 3. **(A)** Immunohistochemical labeling of larval *Tg(isl2b:tagRFP)* retina sections show that RGCs, located in the innermost ganglion cell layer (GCL) of the retina, are robustly and uniformly labeled by RFP. Scale bar, 50 µm. **(B)** 3D projection of RGC axons labeled in a live *Tg(isl2b:tagRFP)* larva. Anatomical locations corresponding to arborization fields (AF) 4, 7 and 9 and the tectum are marked. D, dorsal; P, posterior; M, medial. Scale bar, 50 µm. **(C)** Confocal plane across retinotectal layers in a live *Tg(isl2b:tagRFP)* larva shows complete labeling of all innervation domains: SO, stratum opticum; SFGS, stratum fibrosum et griseum superficiale; SGC, stratum griseum centrale; SAC/SPV, stratum album centrale/stratum periventriculare. A, anterior; M, medial. Scale bar, 50 µm. **(D)** Barplot showing relative proportions (y-axis) of RGCs and non-RGC contaminant classes in replicates of larval samples (x-axis). Cell numbers from each replicate are indicated on top. **(E)** tSNE visualization of all larval RGC clusters combined with cells colored by sample of origin, showing lack of strong batch effects. **(F)** tSNE visualization of 29 transcriptional clusters (colors) derived from 11,406 captured larval transcriptomes showing both immature and mature RGCs (points). Clusters are ordered in decreasing relative frequency. **(G)** Violin plots showing the distribution of the number of transcripts detected per cell (y-axis) in each larval RGC cluster (x-axis). **(H)** Violin plot showing the distribution of the number of genes detected per cell (y-axis) in each larval RGC cluster (x-axis). **(I)** Relative frequency (y-axis) of larval RGC clusters (x-axis), ordered from highest to lowest. **(J)** Dotplot showing the top three selectively enriched markers (columns) in each larval RGC cluster (rows). Representation and row ordering as in **Figure 3A**.

**Figure S3.**
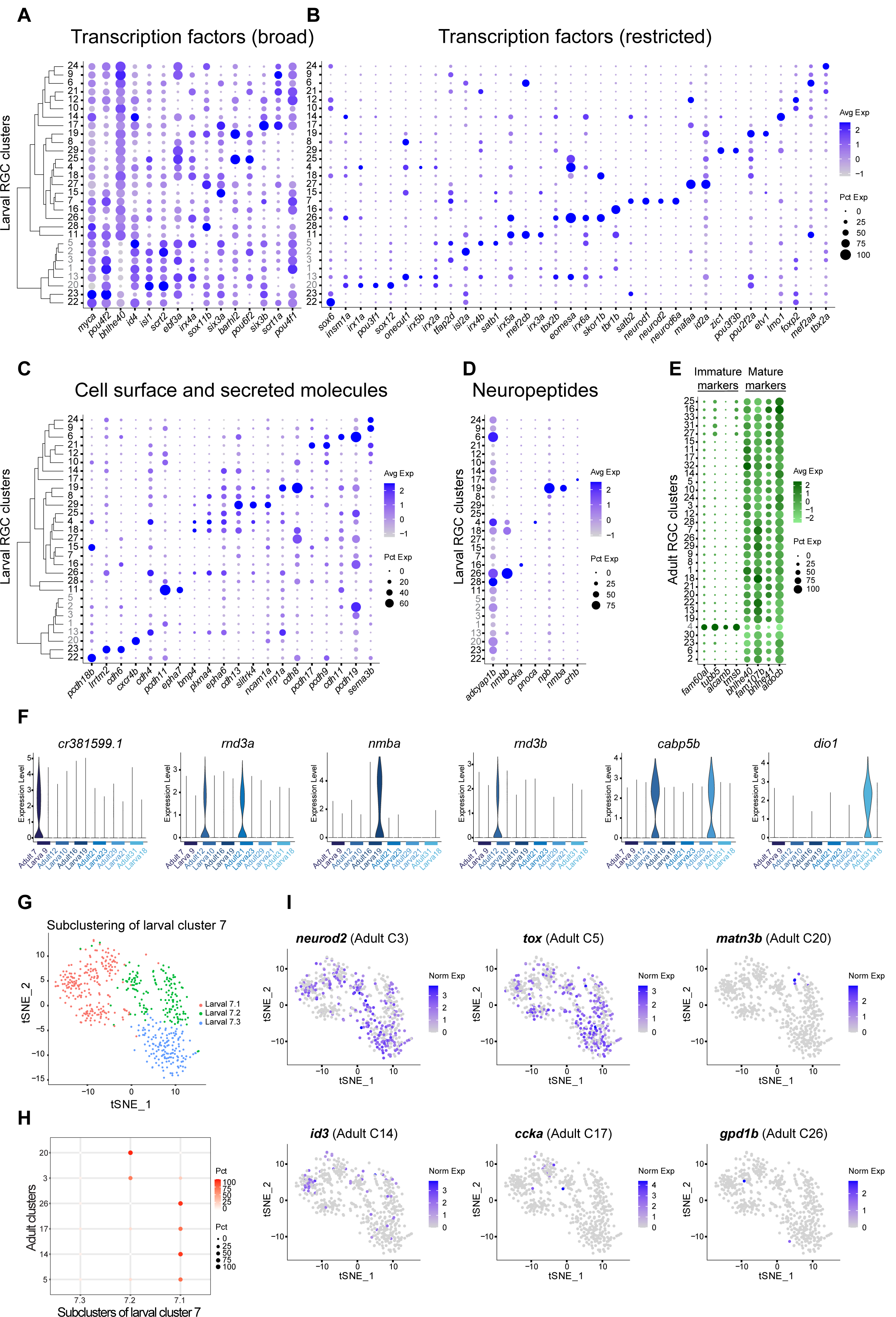
Variably expressed genes across larval RGC types and maturational changes, related to Figure 3. **(A-B)** Dotplots highlighting variably expressed TFs in larval RGC clusters, subdivided into broad (A), and restricted (B) categories. Representation as in **Figure 3A**. The full list is provided in **Table S2**. **(C)** Dotplot highlighting key cell surface and secreted molecules selectively expressed in larval RGC clusters. **(D)** Dotplot highlighting neuropeptides selectively expressed in larval RGC clusters. **(E)** Dotplot highlighting genes (columns) that distinguish immature versus mature larval RGC clusters (as in **Figure 3A**) across adult RGC clusters (rows), ordered as in **Figure 1G**. **(F)** Violin plots showing genes that are up- or down-regulated during maturation in a type-specific manner. Expression values (y-axis) of genes (panels) are plotted for each of the six 1:1 matching larval and adult clusters in panel D (columns). Colored bars indicate matching cluster pairs. **(G)** tSNE visualization of 3 subclusters identified within larval cluster 7 (7.1-7.3). Subclusters were determined through a separate analysis of larval cluster 7 cells (n=539). **(H)** Transcriptional correspondence between adult clusters that map to larval cluster 7 in **Figure 3D**, and subclusters as in panel F, summarized as a confusion matrix. Circles and colors indicate the proportion of cells in an adult cluster (row) mapped to a corresponding subcluster (column) by a supervised classifier trained on subclusters within larval cluster 7. **(I)** Feature plots of genes that are selectively enriched in adult clusters that map to larval cluster 7, reported in **Figure 1H**. Cells within larval cluster 7 (points) visualized as in panel F are colored based on their expression of selected genes (panels).

**Figure S4.**
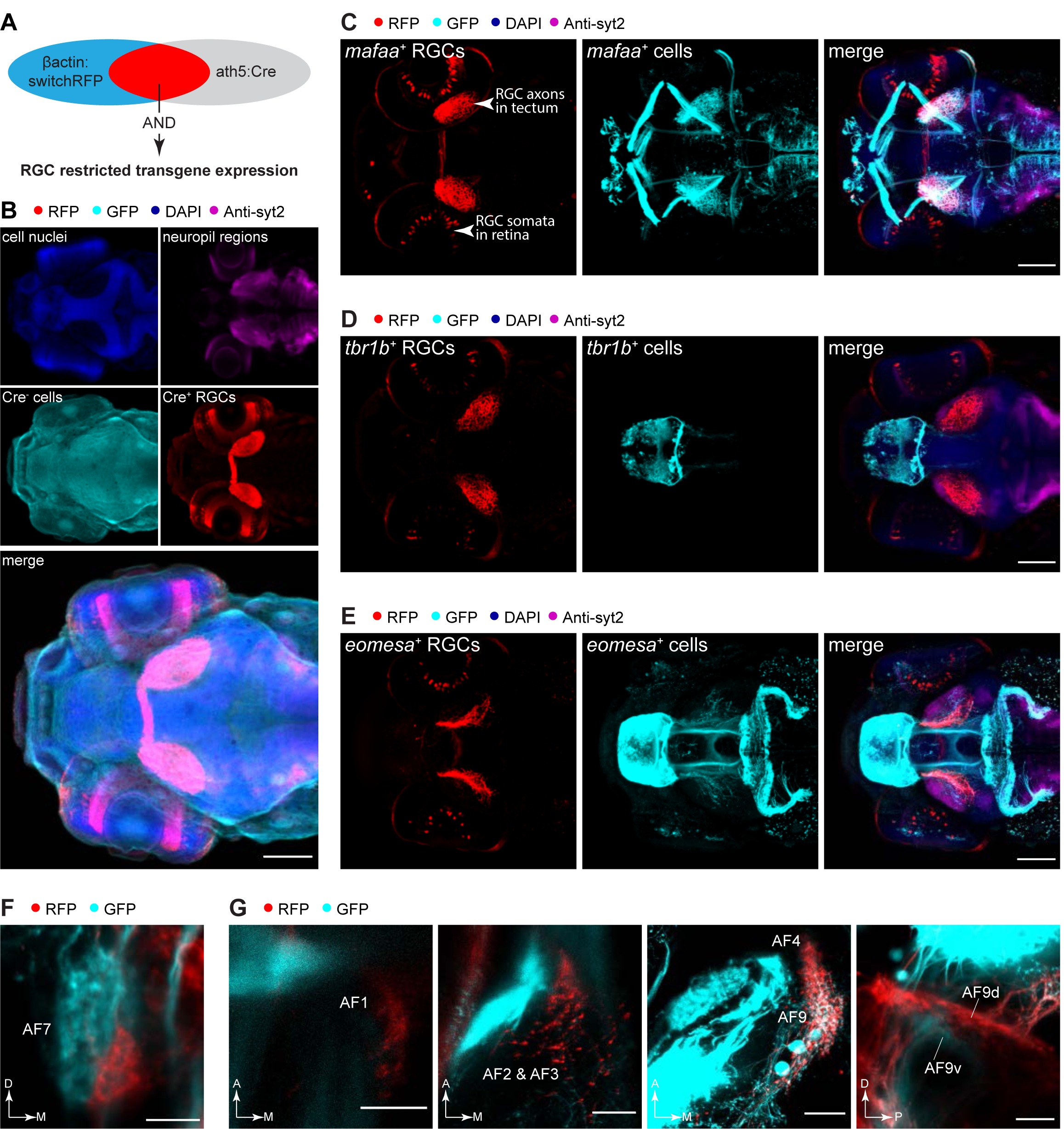
Distinct anatomical features of molecularly defined RGC subclasses, related to Figure 4. **(A)** Genetic intersection via “AND” logic using the RGC-specific line *Tg(ath5:Cre)* refines transgene expression to RGCs in a ubiquitous driver line. **(B)** Immunostained *Tg(ath5:Cre, ß-actin:loxP-GFP-loxP-tagRFP)* larvae show that Cre drives recombination specifically in RGCs as shown by GFP to RFP conversion. Cre^-^ cells continue to express the default reporter GFP. Scale bar, 100 µm. **(C-E)** Characterization of novel RGC-cluster specific driver lines. Overview of immunostained *Tg(driver:QF2, QUAS:switchNTR, ath5:Cre)* larvae, where the “driver” corresponds to *mafaa* (C), *tbr1b* (D), and *eomesa* (E). Besides RFP-labeled cluster-specific RGCs, each marker is expressed in additional tissues: *mafaa* is expressed in muscles and neurons in midbrain and hindbrain, *tbr1b* is expressed in the forebrain and habenula, and *eomesa* is expressed in forebrain, habenula and cerebellum. Scale bar, 100µm. **(F)** Frontal confocal view of AF7 innervated by *mafaa*^*+*^ RGCs in the ventromedial neuropil as labeled in live *Tg(mafaa:QF2, QUAS:switchNTR, ath5Cre, isl2b:GFP)* larval fish. D, dorsal; M, medial. Scale bar, 10 µm. **(G)** Confocal images of divergent extra-tectal *eomesa*^*+*^ RGC innervations in AF1, AF2, AF3, AF4 and pretectal dorsal AF9 (AF9d) but not ventral AF9 (AF9v). A, anterior; M, medial; D, dorsal; P, posterior. Scale bar, 20 µm.

**Figure S5.**
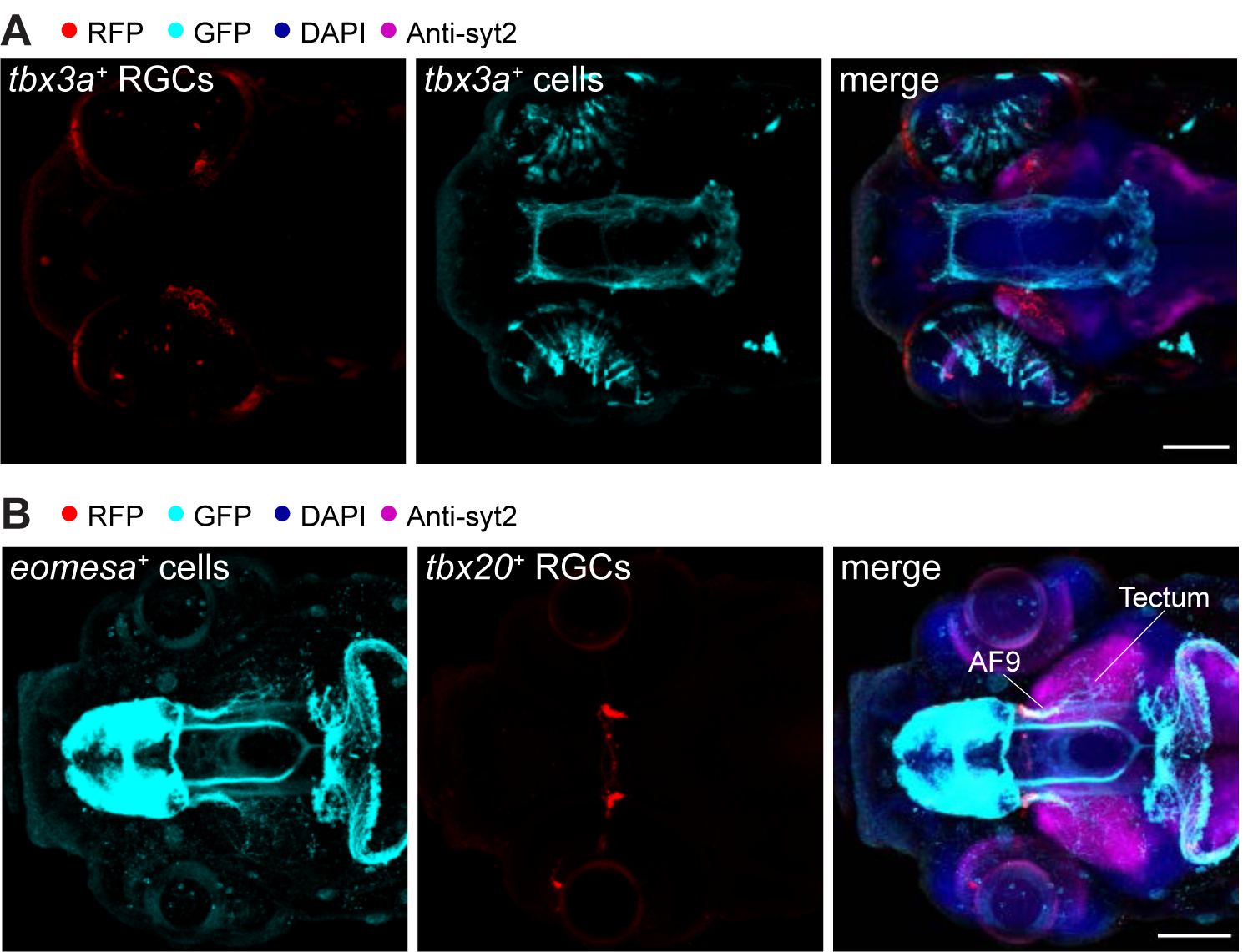
Unique morphotypes within RGC subclasses, Related to Figure 5. **(A)** Overview of immunostained *Tg(tbx3a:QF2, QUAS:switchNTR, ath5:Cre)* larvae. Besides RFP-labeled RGC types, *tbx3a* is expressed in other retinal cell types (Müller glia and bipolar cells) and hypothalamic neurons. Scale bar, 100 µm. **(B)** Immunostained *Tg(eomesa:QF2, QUAS:GFP, tbx20:Gal4, UAS:NTR-mCherry)* larvae. Because there is no crosstalk between the binary expression systems Gal4-UAS and Q-system, *eomesa*^*+*^ cells and *tbx20*^*+*^ cells could be labeled independently and co-visualized in GFP and RFP, respectively. Scale bar, 100 µm.

**Figure S6.**
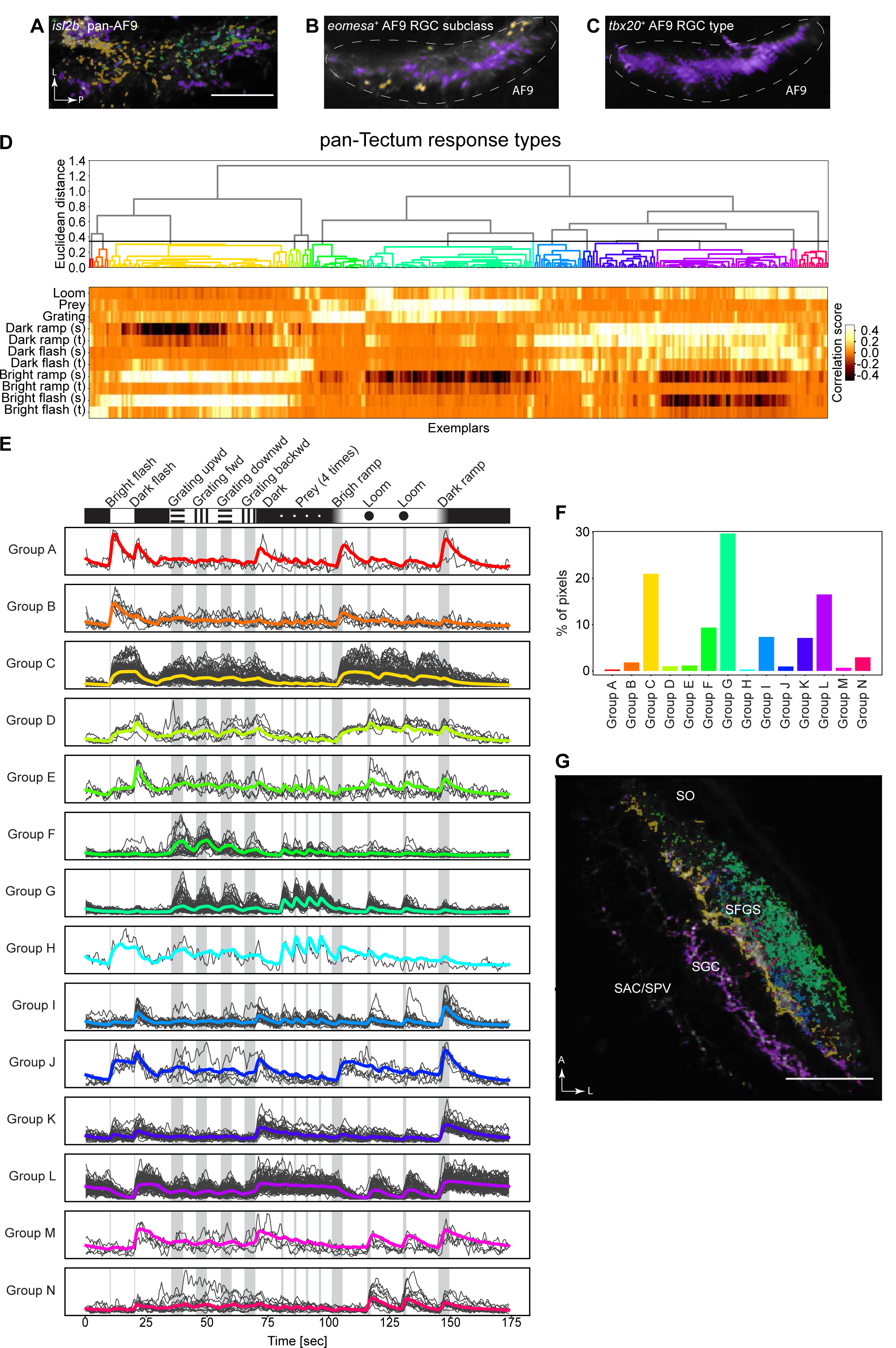
Distinct responses across retinotectal laminae, Related to Figure 6. **(A-C)** Representative recorded neuropil with pixels color-coded by response group assignment as determined in **Figure 6G-I** for *isl2b*^*+*^ RGCs (A), *eomesa*^*+*^ RGCs (B) and *tbx20*^*+*^ RGCs (C). L, lateral; P, posterior. Scale bar, 20 µm. **(D)** Fourteen main response groups can be distinguished from functional imaging across retinotectal laminae using *Tg(isl2b:Gal4, UAS:GCaMP6s)* larvae. Shown is the dendrogram from hierarchical clustering of response exemplars. Neuronal activity to a certain type of visual stimulus is indicated by the correlation score. **(E)** Activity traces of fourteen classified tectal response groups to the stimulus sequence. Shown are the average traces (colored) and all representing exemplars that fall into the group (grey). **(F)** Quantification of relative frequency of *isl2b*^*+*^ RGCs response groups in the tectum. **(G)** Pixels originating from distinct RGC axons innervating tectal laminae were color coded by response group assignment. A, anterior; L, lateral. Scale bar, 50 µm.

**Figure S7.**
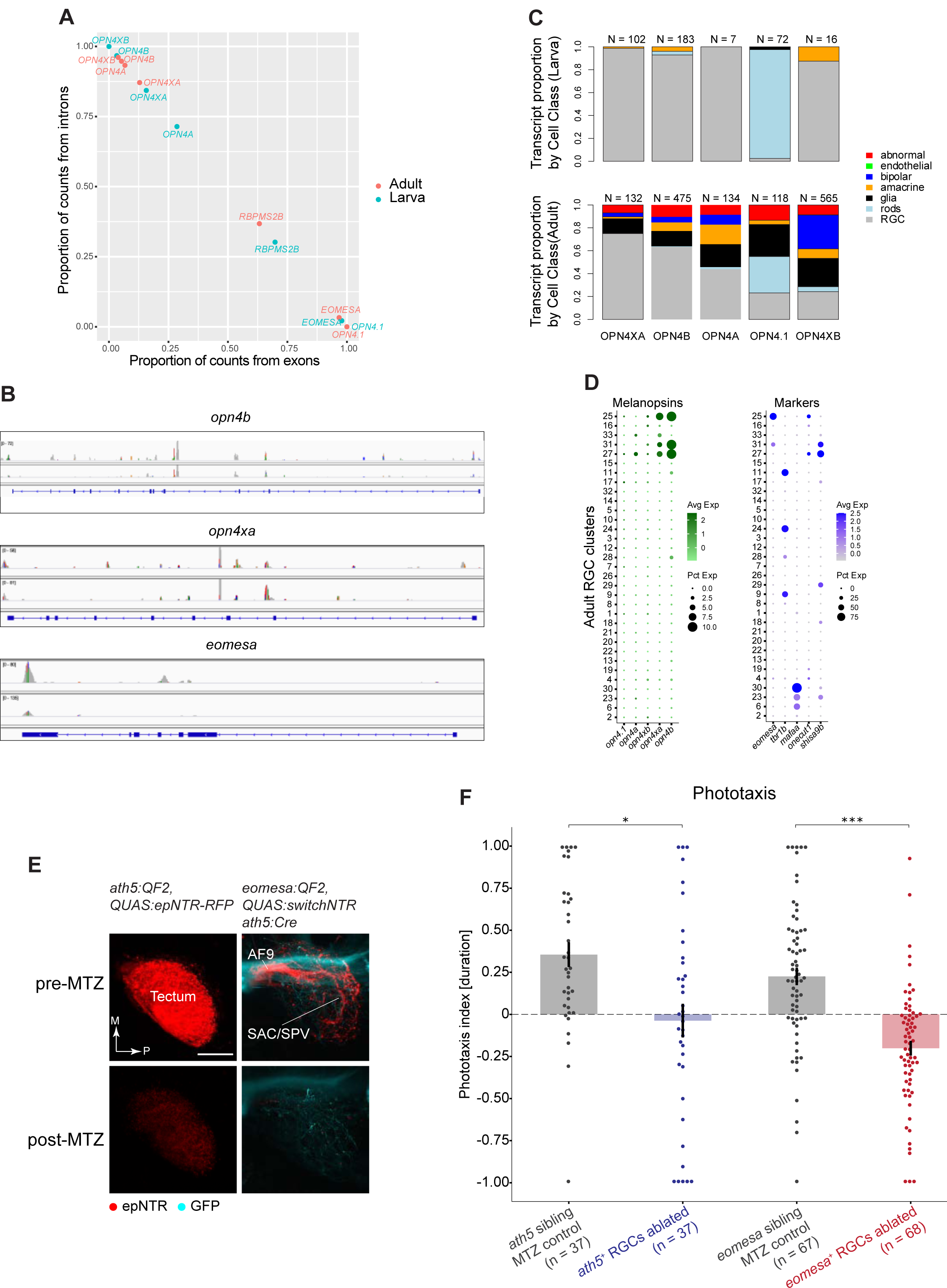
Intrinsic photosensitivity in the zebrafish retina and its implication in phototaxis, Related to Figure 7. **(A)** Relative proportions of intron (y-axis) and exon (x-axis) annotated transcripts for melanopsin genes, *eomesa*, and *rbpms2b* in the larval (blue) and adult (red) data. These counts were computed using velocyto (La Manno et al., 2018). **(B)** Read alignment patterns at the loci corresponding to *opn4b* (*top*), *opn4xa (middle)* and *eomesa (bottom)*. For each gene (panel), the gene body definition is provided in the bottom-most row, with exons denoted by boxes, introns by lines and the arrows indicating the 5’ to 3’ direction. The pile-up of read counts from two separate adult samples shown for each gene. While peaks for *eomesa* correspond to exonic locations, those for *opn4b* and *opn4a* are predominantly derived from introns, consistent with panel A. **(C)** Barplots showing relative frequency (y-axis) of detection of transcripts of melanopsin genes (x-axis) in various cell classes in the larval (top) and adult (bottom) dataset. Cell classes, denoted by colors, correspond to those in **Figures S1A** and **S2D**, respectively. Recovered numbers of transcripts (intronic + exonic) are indicated on the top of each bar for each gene. **(D)** Dotplots showing type-specific expression of melanopsin genes in adult RGCs, as in **Figure 7A**. *Left*: Of the five melanopsin homologs (columns), only *opn4xa* and *opn4b* have discernible expression in specific adult RGC clusters (rows) when intronic reads are accounted for. *Right: opn4xa* and *opn4b* expressing clusters include *eomesa*^*+*^ RGC types but not *mafaa*^*+*^ or *tbr1b*^*+*^ types. C27, the only adult *eomesa*^*-*^ RGC type that expresses melanopsin, is marked by the co-expression of *onecut1* and *shisa9b*. **(E)** Maximum projections of the tectum of live imaged transgenic fish expressing nitroreductase (NTR) in *ath5*^*+*^ RGCs (left) and *eomesa*^*+*^ RGCs (right) before and after treatment with metronidazole (MTZ). M, medial; P, posterior. Scale bar, 50 µm. **(F)** Phototaxis index (PI) values as determined by time spent (duration) in the dim or lit half of the arena for all tested groups: NTR^-^ *Tg(ath5:QF2)* and *Tg(eomesa:QF2)* control siblings as well as *ath5*^*+*^ RGC-ablated blind fish and *eomesa*^*+*^ RGC-ablated larvae. Each dot represents one fish. Error bars represent SEM. * p<0.05, *** p<0.001 (Dunn post-hoc test).

**Table S1. Summary of differentially expressed genes in larval and adult RGC clusters, computed using MAST**.

**Table S2. Summary of expression of transcription factors, neural recognition molecules, and neuropeptides expressed selectively in larval and adult RGC clusters**.

**Table S3. gRNA sequences for CRISPR/Cas9-mediated transgenesis**.

**Video S1. Expression pattern of the *mafaa* transgenic line**.

**Video S2. Expression pattern of the *tbr1b* transgenic line**.

**Video S3. Expression pattern of the *eomesa* transgenic line**.

**Video S4. Expression pattern of the *tbx3a* transgenic line**.

